# Single-molecule FRET probes allosteric effects on protein-translocating pore loops of a AAA+ machine

**DOI:** 10.1101/2023.02.02.526786

**Authors:** Marija Iljina, Hisham Mazal, Ashan Dayananda, Zhaocheng Zhang, George Stan, Inbal Riven, Gilad Haran

## Abstract

AAA+ proteins (ATPases associated with various cellular activities) comprise a family of powerful ring-shaped ATP-dependent translocases that carry out numerous vital substrate-remodeling functions. ClpB is a AAA+ protein disaggregation machine that forms a two-tiered hexameric ring, with flexible pore loops protruding into its center and binding to substrate-proteins. It remains unknown whether these pore loops contribute only passively to substrate-protein threading or have a more active role. Recently, we have applied single-molecule FRET (smFRET) spectroscopy to directly measure the dynamics of substrate-binding pore loops in ClpB. We have reported that the three pore loops of ClpB (PL1-3) undergo large-scale fluctuations on the microsecond timescale that are likely to be mechanistically important for disaggregation. Here, using smFRET, we study the allosteric coupling between the pore loops and the two nucleotide binding domains of ClpB (NBD1-2). By mutating the conserved Walker B motifs within the NBDs to abolish ATP hydrolysis, we demonstrate how the nucleotide state of each NBD tunes pore loop dynamics. This effect is surprisingly long-ranged; in particular, PL2 and PL3 respond differentially to a Walker B mutation in either NBD1 or NBD2, as well as to mutations in both. We characterize the conformational dynamics of pore loops and the allosteric paths connecting NBDs to pore loops by molecular dynamics simulations and find that both principal motions and allosteric paths can be altered by changing the ATPase state of ClpB. Remarkably, PL3, which is highly conserved in AAA+ machines, is found to favor an upward conformation when only NBD1 undergoes ATP hydrolysis, but a downward conformation when NBD2 is active. These results explicitly demonstrate a significant long-range allosteric effect of ATP hydrolysis sites on pore-loop dynamics. Pore loops are therefore established as active participants that undergo ATP-dependent conformational changes to translocate substrate proteins through the central pores of AAA+ machines.

**Statement of Significance:** Molecular machines function by coupling the energy of ATP hydrolysis to mechanical motion. How this coupling occurs and what timescales are involved remains an open question. In this study, we use a powerful single-molecule FRET technique to measure the real-time dynamics of pore loops, which are essential protein-translocating elements of the ATP-dependent disaggregation machine ClpB. Using a series of mutations of the ATP-hydrolysis motifs of ClpB, we find that, although the motions of these pore loops take place on the microsecond time scale, they are markedly affected by the much slower changes in the nucleotide state of the machine. Generally, this study shows that protein machines, such as ClpB, are wired to harness ATP binding and hydrolysis to allosterically affect distal events, such as the function-related mechanics of pore-loops.

## Introduction

Members of the AAA+ (ATPases associated with various cellular activities) protein family are diverse molecular machines that perform multiple ATP-dependent biological functions in cells (1,2). In their functional forms, these proteins typically assemble into asymmetric hexameric rings that can translocate client substrate-DNA (3,4) or substrate-proteins through their central pore (5,6). The residues that constitute ATPase pockets, located at the protomer interfaces (Fig. 1 a), are the highly conserved Walker A motif, GXXXXGK[T/S] (where X is any amino acid), involved in ATP binding, and the Walker B motif, hhhD[D/E] (where h is a hydrophobic residue), essential for ATP hydrolysis (7). The substrate-proteins are bound by flexible pore loops that protrude into the central pore (2). ClpB is a bacterial AAA+ disaggregase that has two nucleotide-binding domains, NBD1 and NBD2, within each protomer (8) and forms a two-tiered hexameric ring (9). Like other members of the AAA+ family, it binds its substrate-proteins by a set of pore loops lining the central pore, including pore loop 1 and pore loop 2 (PL1 and PL2) located in NBD1 and pore loop 3 (PL3), located in NBD2 (9,10) (Fig. 1 a,b).

**Figure 1.**
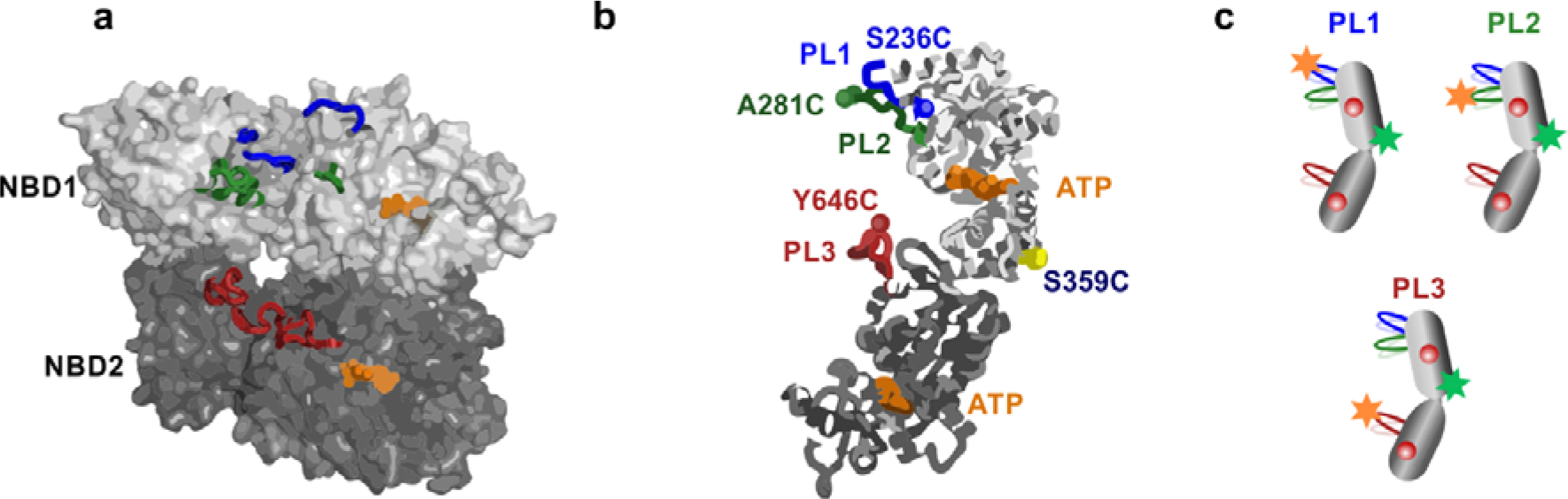
Location of pore loops in ClpB hexamer and fluorescence labeling scheme for single-molecule experiments. **(a)** Side-view of ClpB hexamer from *E. coli* (PDB-6OAX) (9), with three protomers removed to reveal the central pore. The topmost N-terminal domain is absent in this structure. Nucleotides at NBD1 and NBD2, bound at the subunit interfaces, are shown in orange. PL1, PL2 and PL3 of the three shown protomers (A, B and F) are colored in blue, green and red, respectively. **(b)** Monomer structure of ClpB (PDB-6OAX, protomer A (9)). Pore loop 1 (PL1) residues 235-245, corresponding to the sequence GSLLAGA**KYRG**, is shown in blue. Pore loop 2 (PL2) residues 272-290 (LHTVV**GAG**KAEGAVDAGNM) is in green. Pore loop 3 (PL3) residues 637-650 (IGAPP**GYVG**YEEGG) is in red. Numbering and the primary sequence are as in full-length *Thermus thermophilus (TT)* ClpB, used in all experiments in this work. Conserved residues of the primary sequence are in bold. (A short additional pore loop adjacent to PL3, identified in the PDB-6OAX (9) structure, was found to be too close to PL3 and was not studied here.) Residues S236C, A281C and Y646C, used for fluorescence labeling of the pore loops, are shown as spheres on PL1, PL2 and PL3, while residue S359C, used as a reference position in FRET assays, is shown in yellow. Residue numbers are from *TT* ClpB. **(c)** Cartoon representation of the fluorescence double-labeling scheme to study PLs, showing the positions of two Alexa Fluor dyes (as stars) within ClpB monomer. Red spheres represent ATP bound to the two NBDs of ClpB. PL1 construct S236C-S359C, PL2 construct A281C-S359C and PL3 construct S359C-Y646C. In all experiments, we use N-terminally truncated *TT* ClpB.

The ATP-dependent substrate-protein translocation across the central pore of AAA+ machines is thought to be driven by large-scale rearrangements of individual protomers within the hexameric rings (1,2), although the precise mechanism still remains debatable. Multiple recent cryo-EM reconstructions of substrate-bound AAA+ hexamers suggested a structure in which the protomers within the rings are mobile (2) and generally show a spiral arrangement of protomers. Consistent with these structures, substrate-protein translocation is proposed to occur following sequential ATP hydrolysis in a ‘hand-over-hand’ mechanism. In this mechanism, upon ATP hydrolysis and re-binding, subunits within a hexamer move one-by-one, resulting in a unidirectional translocation of the substrate-protein across the central pore with a uniform step size of two residues (3,4). This mechanism is proposed to describe the operation of increasingly more complex AAA+ systems (11). However, the sequential ATP hydrolysis and its associated hand-over-hand translocation mechanism might not be applicable to all AAA+ machines, and other models, concerted and probabilistic ATP hydrolysis, were also put forward for several systems. For example, structural analysis of the hexameric helicase LTag suggested concerted nucleotide binding and exchange mechanism (12). Furthermore, two relatively recent cryo-EM studies of substrate-bound ClpXP yielded highly similar hexameric staircase structures but two different interpretations were offered, and either a sequential (13) or a probabilistic mechanism was proposed (14,15). Biochemical studies on covalently tethered ClpX (16) and HslU (17) found that these AAA+ hexamers remained functional even with multiple inactive subunits, supporting a probabilistic mechanism. In addition, single-molecule optical tweezers assays with ClpXP (18) and ClpB (19), and high-speed AFM experiments with Abo1 (20) yielded results that were more consistent with their probabilistic activities.

Pore loops lining the central channel of AAA+ proteins are essential substrate-binding elements, and their mutations severely impair translocation activities of these machines (2). Pore loop 1 and pore loop 3 in ClpB contain conserved residues KYRG and GYVG, respectively (8) (Fig. 1 b). The highly conserved functional tyrosines are well-known to bind to client proteins in ClpB (6,21) and in other AAA+ machines (22,23). Furthermore, structural analyses revealed that in these pore loops, flanking hydrophobic amino acids are also in contact with substrate-proteins (10,14). Not only the primary sequence, but also the spatial arrangement of the substrate-binding pore loops within AAA+ hexamers is remarkably well conserved.

They form an almost identical spiral-staircase pattern around the bound substrate in a vast number of cryo-EM structures of AAA+ hexamers including Yme1, spastin, proteasome, Cdc48, NSF amongst many others (2). The primary sequence of pore loop 2 is not well-conserved, although its residues were also found to be in contact with protein-substrates in ClpB (9,24) and in its yeast analogue Hsp104 (10). Furthermore, its func tional significance for disaggregation was verified by mutational analysis (10).

Interestingly, increasing experimental evidence indicates that conformational changes of the substrate-binding pore loops contribute to the ATP-dependent translocation by AAA+ machines. Indeed, based on monitoring fluorescence from a tryptophan mutant of pore loop 1 in ClpB, it was suggested that this pore loop undergoes conformational changes that depend on the type of bound nucleotide (21). Furthermore, based on disulfide-crosslinking of substrate-peptides to pore loops in ClpX (25) and on mutational and functional analyses (26), it was proposed that the pore loops undergo ATP-dependent structural changes between ‘up’ to ‘down’ conformations along the axial channel to propel the bound substrate-protein through the hexameric ring. These bulk biochemical studies could not yield a structural model to describe the detected conformational changes. More recently, however, several structural studies explicitly visualized distinct conformational states of the pore loops. From analysis of the structures, it could be inferred that the GYVG pore loops in ClpB and ClpXP respond to substrate-protein binding (13,24). Moreover, pore loop 2 was captured in distinct ‘up’ and ‘down’ conformations in the crystal structure of ClpB’s yeast analogue Hsp104 (10).

While recent advances in structural methods have provided unprecedented details on substrate-protein binding by AAA+ proteins and even on the conformational changes within their pore loops, the extent and timescales of these motions, and their potential coupling to the ATPase activity of these proteins remain to be characterized. In particular, the relationship between the nucleotide state of individual protomers and the conformations of their pore loops are challenging to study due to averaging of the nucleotide-binding sites in cryo-EM (2). Moreover, detecting the real-time dynamics of distinct pore loop types during the ATP-dependent activity of AAA+ proteins under native conditions in aqueous solution requires suitable experimental techniques. Recently, we employed a powerful combination of single-molecule FRET (smFRET) spectroscopy and photon-by-photon Hidden Markov modeling (27) to study the dynamics of pore loops in ClpB under conditions of active ATP hydrolysis (28). We found that these pore loops fluctuate on the microsecond time scale between two major conformations, ‘up’ and ‘down’ along the central pore and that the populations of these two conformations change upon binding to substrate-protein. Furthermore, we found that the dynamics of pore loops 2 and 3 respond to ATP hydrolysis, and the dynamics of pore loops 1 and 3 are correlated to the bulk disaggregation activity of ClpB. These differential responses led us to propose that the pore loops act as Brownian ratchets to translocate substrate-proteins through the central channel. In particular, we suggested that the ATP-dependent pore loops 2 and 3 both serve as pawls that rectify substrate protein translocation through ClpB’s central channel. Here, we set for the first time to investigate the relationship between the nucleotide state of each of the two NBDs of a ClpB hexamer and pore-loop dynamics. To this end, we introduce mutations into the conserved Walker A and Walker B motifs in either NBD1 or NBD2 or both in order to restrict ClpB to pre-defined ATP-activity states, and characterize these constructs by smFRET spectroscopy. Furthermore, we study the potential coupling between the NBDs and the pore loops by molecular dynamics simulations to gain residue-level insights. We demonstrate that the nucleotide states of NBD1 and NBD2 separately can alter the dynamics of all three sets of pore loops, and pore loops 2 and 3 experience the most prominent modulations. Thus, our results reveal unexpected cooperative allosteric interactions between both ATP binding sites and the pore loops of ClpB, and shed light on how ATP binding and hydrolysis drive substrate-protein translocation by the machine.

## Materials and Methods

### Protein expression and purification

All mutant proteins were generated through standard site-directed mutagenesis. *Thermus thermophilus* ClpB variant with truncated N-terminal domain, starting at residue 141 (Val), (referred to as “ClpB” throughout), and its mutants, cloned into a pET28b vector, were expressed in *E. coli* and purified as described previously (28,29). The truncated ClpB was selected to avoid any hindrance to the fluorescent dye on PL1 by the N-domain, and was previously verified to be fully assembled and functional (28).

### Protein labeling and protomer mixing

Double-cysteine mutants of ClpB were labeled with Alexa Fluor 488 C5 maleimide (AF488) and Alexa Fluor 594 C5 maleimide (AF594) dyes (Thermo Fisher) following a protocol described in our previous work (28). Any unreacted dye molecules were removed on a desalting column (Sephadex G25, GE Healthcare). For the preparation of ClpB hexamers with only one labeled subunit, suitable for smFRET measurements, AF488-AF594-labeled double-cysteine mutants were combined with 100-fold molar excess of unlabeled cysteine-less ClpB. With this ratio, the probability for the incorporation of one labeled protomer in a hexamer was 5.7%, whereas the probability to find two labeled protomers in the same hexamer was as low as 0.15%. To achieve full refolding and homogeneous reassembly, the protein solutions were initially dialyzed in the presence of 6 M GdmCl. This was followed by dialysis steps in the presence of 4 M, 2 M, 1 M and 0 M GdmCl. The final steps involved extensive dialysis into low-salt buffer (25 mM HEPES, 25 mM KCl, 10 mM MgCl_2_, 2 mM ATP, pH 8) and filtration through 0.1 μm filters (Whatman Anotop-10). The assembled ClpB was aliquoted, flash-frozen and stored at -80°C until further use. For the preparation of hexamers containing a functional mutation (any of the Walker mutants described in this study), fluorescently double-labeled ClpB bearing the mutation was mixed with the 100-fold excess of unlabeled cysteine-less ClpB containing the same functional mutation, and reassembled following the above procedure.

### ATPase activity measurements

ATP activity of ClpB mutants was measured using a coupled colorimetic assay (30). ClpB or its mutants (1 μM total monomer concentration) were incubated in the presence of 2 mM ATP in 50 mM HEPES (pH 8), 50 mM KCl and 0.01% Tween 20, with an ATP regeneration system (2.5 mM phosphoenol pyruvate, 10 units/ml pyruvate kinase, 15 units/ml lactate dehydrogenase, 2 mM 1,4 dithioerythritol, 2 mM EDTA and 0.25 mM NADH). To assess the effect of the model substrate k-casein (Sigma Aldrich), it was added to a final concentration of 25 μM. ATP hydrolysis was initiated by adding MgCl_2_ (10 mM), and measured by monitoring the decrease in NADH absorption over time at 340 nm, using a microplate reader (Synergy HTX, BioTek) equilibrated at 25°C. The rate of ATP hydrolysis was determined from the initial linear slope of the measured data. ATP hydrolysis rate per ClpB monomer per minute is reported.

### smFRET measurements

Custom-made glass flow chambers for smFRET measurements were prepared as previously reported (31). The chambers were coated with a supported lipid bilayer composed of egg phosphatidylcholine (Avanti Polar Lipids) to prevent protein absorption to the glass surface. The reassembled hexamers of ClpB were diluted to ∼50 pM of labeled ClpB, which corresponds to ∼5 nM of total ClpB, into buffer (25 mM HEPES, 25 mM KCl, 10 mM MgCl_2_, 2 mM ATP, 0.01% Tween 20, pH 8), loaded into the chambers and sealed with silicon grease. Experiments in the absence of ATP and Mg^2+^ (Fig. S3) were performed by simply omitting these chemicals from the solution; it is possible that residual ATP/Mg^2+^ might still remain in these samples from the protein purification procedures, but their quantities should be too small to affect the proteins. Measurements on freely diffusing molecules were conducted as described before (28), using a MicroTime 200 confocal fluorescence microscope (PicoQuant). The samples were excited using a pulsed interleaved excitation scheme with 485 nm and 594 nm diode lasers pulsed at a 3:1 ratio, with the repetition rate of 40 MHz, operating at 50 μW and 8 μW, respectively. The emitted photons were split into two channels by a dichroic mirror (FF580-FDi01, Semrock), and passed through band-pass filters (520/35 nm, BrightLine, Semrock, for the AF488 emission and ET-645/75m, Chroma, for the AF594 emission). Photon arrival times were detected by two single-photon avalanche photo-diodes (Excelitas SPCM-AQR-14-TR) coupled to a standalone time-correlated single photon counting module (HydraHarp 400, PicoQuant). Data were acquired for around 5 h per sample at a fixed ambient temperature (22°C), and no evidence of any temperature or heating effects was observed. At least two samples were analyzed per condition.

### smFRET data analysis and H^2^MM analysis

Fluorescence bursts, corresponding to the single molecules of ClpB, were selected using data analysis workflows developed in the lab (27-29,31). A cut-off of 5 μs was used to effectively separate fluorescence bursts from the background. Raw FRET efficiency and raw stoichiometry values were calculated as described (32), and a two-dimensional histogram of raw stoichiometry against raw FRET efficiency was constructed and used to calculate correction factors, the leak (∼0.05) and the direct excitation (∼0.02). Following data correction using these factors, FRET efficiency for each burst was calculated (as photons arriving from the acceptor channel divided by the total number of photons). To obtain the final corrected FRET histogram without the donor-only and acceptor-only populations, we selected only photon bursts with a stoichiometry corresponding to molecules bearing both active donor and acceptor dyes and containing at least 30 photons. The same parameters for the selection of double-labeled molecules were used throughout. These stringent selection criteria eliminated any potential photobleaching/blinking events. Once the double-labeled molecules were selected, normalized FRET efficiency distribution functions were displayed as histograms with 35 bins. To point out, FRET efficiency distributions were used ONLY for qualitative comparison.

For the quantitative kinetic analysis, we used the same selected data for a maximum likelihood Hidden Markov modeling analysis, H^2^MM, introduced and described in detail previously (27). In this analysis, the arrival time and the type (donor or acceptor) of the selected photons represent the observation sequence (O). The model (λ) comprises i) the probability matrix (Π), ii) the transition matrix (A) and iii) the observation matrix (B). The algorithm performs the following steps: a) initialization, when initial model parameters Π, A and B are guessed (using 50 initial guesses). b) Expectation, when the model parameters Π, A and B are learned given the observation sequence O. This is iterative step and proceeds using the forward-backward Baum-Welch algorithm (a special case of the Expectation-Maximization algorithm. c) Maximization, where Π, A and B are re-calculated based on estimators derived from the previous step. d) Determination of the model’s likelihood, P(O|λ), defined as the conditional probability of observing the sequence O given the model λ. e) Viterbi algorithm: using the best model (λ =(Π, A, B), which is characterized with the highest likelihood across all 50 guesses, the algorithm calculates the most probable sequence of states. For this analysis, we chose ∼7,000 photon trajectories per sample, and analyzed them with a fixed two-state model, where the number and FRET efficiency of the states are fixed (detailed in Supplemental Information, section “Supplemental Data Analysis Details” and Table S1), and other parameters are freely and independently optimized. The choice of the two-state model was based on our preceding finding that the free energy profiles derived from pore-loop data displayed two minima (28). The approach was validated by recoloring and segmentation analyses, as well as dwell time calculations (detailed in Supplemental Information, section “Supplemental Data Analysis Details” and Supplemental Fig. S2 and Tables S6). Effective equilibrium coefficient, defined as 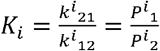 (where *k*^*i*^, are the transition rates and *P*^*i*^ are the populations of state 1 and state 2 for each pore loop) was derived from the H^2^MM analysis of each pore loop type (*i*) Tables S6).

### Molecular dynamics simulations

We performed molecular dynamics (MD) simulations of wild-type and double Walker B (BB) mutant of the *Escherichia coli* ClpB using the Gromacs 2022 package (33) and the GROMOS96 54A7 force field. ClpB structures, truncated at N-terminal domains (amino acids 1-160) and middle domains (amino acids 409-524), were modeled using the cryo-EM structure corresponding to the pre-hydrolysis configuration (PDB-6OAX (9)). We used the Modeller software version 9.23 (34) to build missing loop residues of the pre-hydrolysis conformer, namely amino acid residues 525-528 in protomer 5 and 284-293 in protomer 6 (using protomer numbering in accord with the cryo-EM studies (9)). To ensure chain connectivity in the absence of middle domains, in each chain, residues 408 and 525 were connected through a linker comprising five Gly residues. In the pre-hydrolysis state, protomer 1 is bound to ATPγS in NBD1 and to ADP in NBD2, protomers 2 through 5 are bound to ATPγS in both NBD1 and NBD2, and protomer 6 is bound to ADP in both NBD1 and NBD2. In our simulations, ATPγS and ADP molecules were modeled by using the Automated Topology Builder server that generates force field parameters compatible with the GROMOS96 54A7 force field (35). Wild-type ClpB simulations used the same setup and extended the time scales probed in our previous studies (36). ClpB mutants were modeled by using PyMOL (35) to implement Walker B mutations in the cryo-EM structure, E279A in NBD1 and E678A in NBD2. These MD simulations, principal component analyses, and analyses of optimal and suboptimal paths are described in detail in the Supplemental Information (section “Supplemental Computational Methods”).

### Double-mutant cycles for Walker B mutants

The double mutant cycles (DMCs) were constructed as previously described (37). More specifically, the free energy, Δ*G*_*i*_ in Jmol^-1^, for each PL mutant was calculated from equilibrium coefficients according to:

Δ*G*_*i*_ = -*RTln*(*K*_*i*_) where R=8.314 Jmol^-1^K^-1^ and T=295.15 K. Subsequently, the free energy change associated with a mutation (from WT to B1 or from B2 to BB) was calculated as follows:

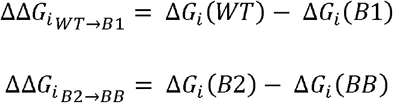

and plotted along the edges of the cycles (Fig. 6d-f and Fig. S15).

In the DMCs analysis, if the ΔΔ*G*_*i*_ values along the opposite edges of the cycle are unequal, that is, if 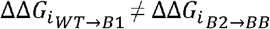 and 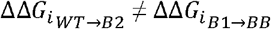 then the residues at the two positions are thermodynamically coupled either through a direct or indirect interaction (37). This is true in our DMCs for all PLs, either with or without κ-casein, indicating that the effects of B1 and B2 mutations are coupled in all cases.

The coupling energy is calculated as

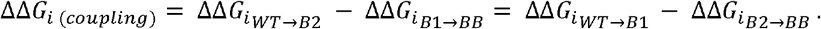

## Results

### Abolishing ATP hydrolysis significantly alters PL2 and PL3, but not PL1 dynamics

To study pore-loop dynamics by smFRET, we labeled ClpB molecules with two fluorescent dyes that comprise a FRET pair, Alexa Fluor 488 and Alexa Fluor 594, as in our preceding study (28). One dye was placed directly onto the pore loop of interest, and the second dye was located on a rigid reference position in the middle of the molecule (Fig. 1 b,c). We previously verified that labeling at these positions does not hinder the activity of ClpB (28). Based on this labeling scheme, it is expected that as the flexible pore loop moves up and down along the axial channel, the relative distance between the two dyes would change, leading to fluctuations in the measured FRET efficiency signal. For smFRET measurements, we mixed the doubly-labeled ClpB protomers with a 100-fold molar excess of unlabeled ClpB protomers, using a refolding and reassembly procedure introduced previously (Materials and Methods). This procedure yields ClpB hexamers that contain a single fluorescently labeled protomer. In case when a functional mutation is present, we always combine the double-labeled ClpB mutant protomer with its corresponding unlabeled cysteine-less ClpB mutant protomers (see Materials and Methods) in order to ensure that the studied functional mutation is present in all protomers of the re-assembled ClpB hexamer, rather than only in the fluorescently labeled protomer. To study the effect of abolished ATP hydrolysis on pore-loop dynamics, we first mutated a conserved glutamic acid residue, acting as an essential water-attacking base for ATP hydrolysis (7), within Walker B motifs in both NBDs (E271A/E668A, denoted as “BB”). To point out, these mutations within the nucleotide-binding sites are distant from the pore loops, as can be seen from the structure of ClpB (Fig. 1). Homogeneous assembly and absence of ATPase activity in these mutants were verified by native PAGE chromatography and ATPase activity assays, respectively (Fig. S1). Pore-loop constructs with (“BB”) or without (“wt”) the mutations were analyzed by smFRET spectroscopy in aqueous solution in the presence of a saturating concentration of ATP (2 mM), either without or with the addition of the soluble model substrate-protein κ-casein (38). We previously estimated that with κ-casein, our results mostly represent the protein-bound protomers (28). Bursts of photons emitted as labeled ClpB molecules freely diffused through a focused laser beam were collected, and FRET efficiency histograms were constructed from the experimental data following selection of double-labeled molecules (Materials and Methods, Fig. 2).

**Figure 2.**
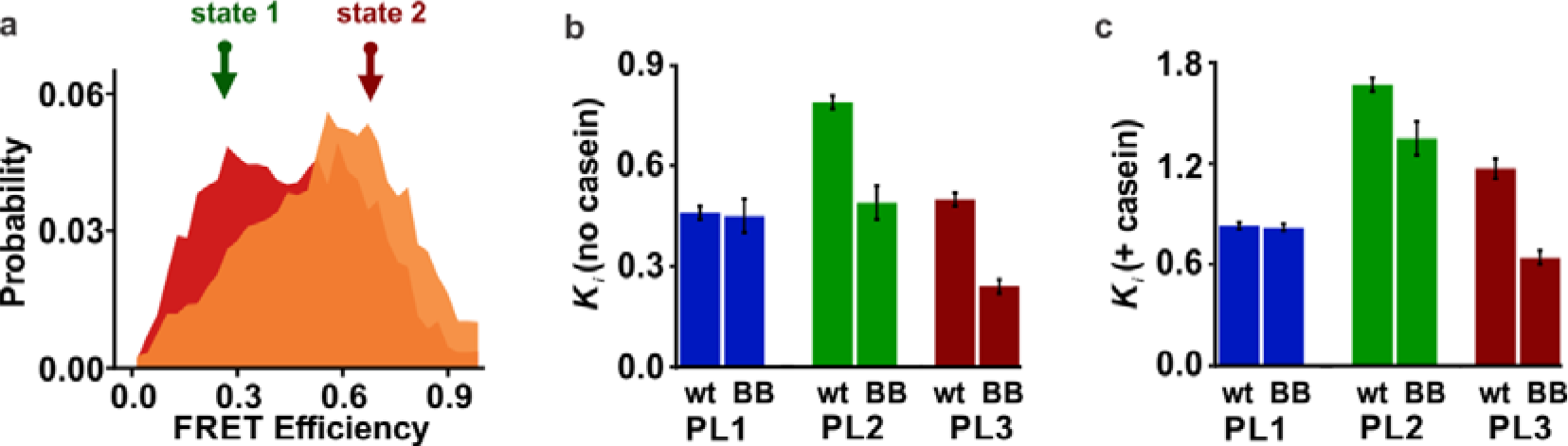
Effect of double-Walker B mutations on the dynamic equilibrium of PL1, PL2 and PL3. **(a)** FRET efficiency histograms of PL3 (red) and PL3 BB (orange), both in the presence of 25 μM κ-casein. Here and elsewhere below, single-molecule measurements were conducted using 1:100 labeled:unlabeled ClpB with 2 mM ATP. BB mutations are E271A/E668A (numbering as in the full-length *TT* ClpB). Arrows show the FRET efficiency values of two fixed states (same across different PL3 mutants) used in the H^2^MM analysis of PL3 (Table S1). The positions of these two states were obtained from a global analysis of pore-loop data with and without κ-casein done previously (28) (detailed in Supplemental Information, under “Supplemental smFRET Data Analysis Details”). See Figure S1 for smFRET histograms of PL1, PL2 and PL3 wt and BB constructs with and without κ-casein. **(b)** H^2^MM-derived equilibrium coefficients, *K*^*i*^s, for wt and BB mutants, measured without κ-casein. **(c)** H^2^MM-derived equilibrium coefficients, *K*_*i*_s, for the same constructs, measured with 25 μM κ-casein. All *K*_*i*_ values are listed in Table S6. Overall higher values than in (b) are due to a clear increase in the population of the low-FRET state in all pore loops in the presence of κ-casein. In both (b) and (c), differential effect of the BB mutation was registered (average values are reported, errors are SD, n=2-3 repeats of the In good agreement with our preceding study (28), the FRET efficiency histograms of the wt pore-loop constructs (without Walker B mutations) were broad, indicating dynamic heterogeneity, and displayed a shift to low FRET efficiency values upon κ-casein addition (Fig. S1). We previously verified that κ-casein addition does not impact the photophysical properties of our fluorescent dyes (as discussed in Supplemental Information, section “Supplemental smFRET Data Analysis Details”). We stress that FRET efficiency histograms in this study are used only for qualitative purposes. To quantitatively characterize the underlying pore-loop dynamics, we analyze fluorescent bursts on a photon-by-photon level, using H^2^MM (27), a statistical tool for the analysis of single-molecule data, in which photon arrival times are used as input (Materials and Methods). As previously (28), we found that these data are best described with two states, state 1 at low FRET efficiency and state 2 at high FRET efficiency, and used fixed FRET efficiency values for the analysis (as detailed in Supplemental Information, section “Supplemental smFRET Data Analysis Details” and Table S1). We assign the two FRET efficiency states to the up and down conformations of the pore loops along the axial channel, based on the H^2^MM analysis and on previous structural triangulation calculations, as well as control measurements that verified the absence of relative motion of the NBDs themselves (28). It is possible that the up conformations of the pore loops serve to engage the substrate-protein and avoid its back-slipping, and the down conformations aid with the directional translocation of the substrate-protein across the axial channel. The H^2^MM analysis was verified by stochastic recoloring of the data and segmentation analysis to confirm the two-state representation (Fig. S2), and by dwell time calculations which were in agreement with the H^2^MM-derived rates (Tables S6). According to the H^2^MM analysis, the transition rates between the two states were fast, 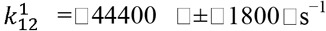 and 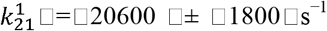 for PL1 without substrate-protein, and slightly slower upon κ-casein addition, 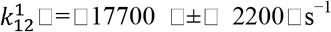 and 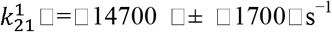 (errors from at least three repeats, see Tables S6 for the transition rates of all other pore-loop types). These microsecond-timescale transition rates position pore-loop motions on a much faster time scale than the ATP hydrolysis in ClpB. To characterize the changes in the state populations of each pore loop type (*i*), we used an effective equilibrium coefficient, 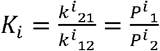, defined as the population ratio of state 1 to that of state 2 (Tables S6) (28). In response to κ-casein addition, the equilibrium coefficient of PL1, *K*_1_, changed from 0.46 ± 0.02 to 0.83 ± 0.02, which was reflected in the shift to low-FRET values in the FRET efficiency histograms and fully agreed with our preceding results (28). PL2 and PL3 both also responded to κ-casein addition by a shift to low-FRET values (Fig. S1). To note, according to our previous analysis (28), our measurements are consistent with pore-loop motions of around 10 Å (or 2 amino acids), representing significant structural fluctuations, which likely involve also adjacent structural elements.

In the BB constructs where ATP is bound and not hydrolyzing, all three PLs still displayed high transition rates between their two states, similar to the non-mutated constructs (Table S6). For example, PL1 BB without κ-casein showed k^*1*^_12_□=□47600 □±□ 6000□s^−1^ and k^*1*^_21_□=□22500 □±□1900□ s^−1^ close to the values for PL1 wt. Furthermore, while the BB mutants still responded to κ-casein addition by a shift to low FRET efficiency values, their responses were modulated in a differential manner (Fig. 2, Fig. S1). We quantified these κ-casein-induced shifts by using parameters derived from the H^2^MM analysis (as summarized in Tables S6). The results corroborate our previous study (28), in which we found that PL1 responds similarly to the substrate in the presence of either ATP or ADP, while PL2 and PL3 responded differently. Importantly, we did not observe this characteristic κ-casein-induced shift in the FRET efficiency histograms under conditions that favor ClpB disassembly, such as high-salt and absence of added nucleotide (Fig. S3). Thus, the κ-casein-induced change in the pore-loop dynamics is a feature of ATP-bound and assembled ClpB complexes. Surprisingly, FRET efficiency histograms of PL3 BB displayed a shift to high FRET efficiency values compared to the corresponding histograms of the non-mutated samples, both with and without κ-casein (Fig. 2a, Fig. S1). Furthermore, the derived H^2^MM parameters for PL2 BB and PL3 BB mutants differed significantly from the results for their corresponding wt constructs, indicating a strong perturbation of the pore-loop dynamics by the BB mutations. These results are summarized in Fig. 2, Tables S6 and Fig. 6 b,c. In particular, the effective equilibrium coefficients were decreased (Fig. 2 and Tables S6). This decrease was especially strong in the case of PL3, with *K*_3_ changing from 0.50 ± 0.02 for PL3 wt to 0.24 ± 0.02 for PL3 BB, and from 1.17 ± 0.06 for PL3 with κ-casein to 0.64 ± 0.04 PL3 BB with κ-casein. This change indicates that the high-FRET state of PL3, which corresponds to the “up” conformation, becomes more populated in the absence of ATP hydrolysis. Considering that the BB mutations are remote from the pore loops, this points to strong allosteric communication between the pore loops and the ATPase sites in ClpB. Therefore, even though pore loops fluctuate on the microsecond-timescale, the dynamic equilibrium of PL2 and PL3 is significantly affected by the changes in the nucleotide state of the machine. In the PL1 samples, however, the difference between the wt and BB samples was absent, indicating that the dynamics of this pore loop are not affected by abolished ATP hydrolysis (28).

### Double-Walker B mutations perturb conformational fluctuations and allosteric paths of PL2 and PL3, but not of PL1

The effect of BB mutations on pore loop motions can be rationalized in terms of the effect of mutations on conformational dynamics of ClpB and the pore loops and on the allosteric networks connecting the ATP-binding sites to the pore loops. To obtain information on dynamics and allosteric networks, we performed molecular dynamics (MD) simulations of wt and BB variants of *Escherichia coli* (*E. coli*) ClpB, for which high-resolution structures are available (9) (see Materials and Methods).

In equilibrium simulations, principal component analysis (PCA) provides significant insight into conformational fluctuations by highlighting the independent modes of motion that collectively determine the essential dynamics. The PCA approach relies on diagonalization of the covariance matrix of atomic fluctuations, which yields eigenvalues that characterize the amplitude of fluctuations and eigenvectors that correspond to the orthogonal directions with maximal variance (see Materials and Methods). We focus on the eigenvectors corresponding to the top eigenvalues with the largest contribution to the variance. We perform this analysis on the pore loops of protomers 2-4, as both NBDs of these protomers are in an active (ATP-bound) state. For PL1 and PL2 pore loops of these protomers, both in the wild-type and BB variant, the top 10 eigenvalues, arranged in decreasing order, contribute > 80% of the variance, therefore we restrict our analysis to the corresponding eigenvectors in all systems considered. Quantitative comparison between the essential subspaces comprising 10 eigenvectors of the wt or BB variants is made by using the root-mean-square inner product (RMSIP) between the two sets of eigenvectors (see Materials and Methods). Intriguingly, pore loops display a distinct response to perturbation. PL1 loops demonstrate a high RMSIP value of ≅ 0.80, which reveals a strong similarity of motions of this set of pore loops and weak perturbation of their dynamics. Consistent with this observation, we note the large overlap between PC1 eigenvectors of PL1 loops in the BB variant and in the wt ClpB, ≅ 0.85. Swing-like motions of PL1 loops observed for the wt ClpB are largely preserved in the BB variant (Figures 3a and Movies SM1). A moderate overlap is found for PC2 eigenvectors of PL1 loops, ≅ 0.54, which emphasize torsional motions in both wt and BB variants (Figures S4a). Motions of PL2 and PL3 loops (Figures 3 b-c and S4 b-c, Movies SM2-3) are more strongly perturbed by the BB mutations than those of PL1 loops, with the RMSIP values of ≅ 0.64 for PL2 and 0.68 for PL3, respectively. Correspondingly, eigenvectors PC1 and PC2 of PL2 and PL3 loops in the BB variant have small to moderate overlap with the top two PCs in the wt ClpB, as indicated by cumulative overlaps of each of the top two PCs in the BB variant with respect to those in wt ClpB ranging between 0.09-0.51 (Table S2). Thus, whereas, for PL1, the two PCs of the BB variant corresponding to the top eigenvalues are well represented by the top two PCs of the wild-type ClpB, for PL2 and PL3 a larger set of PCs is required, corresponding to lower eigenvalues of the wild-type ClpB (Table S2).

**Figure 3.**
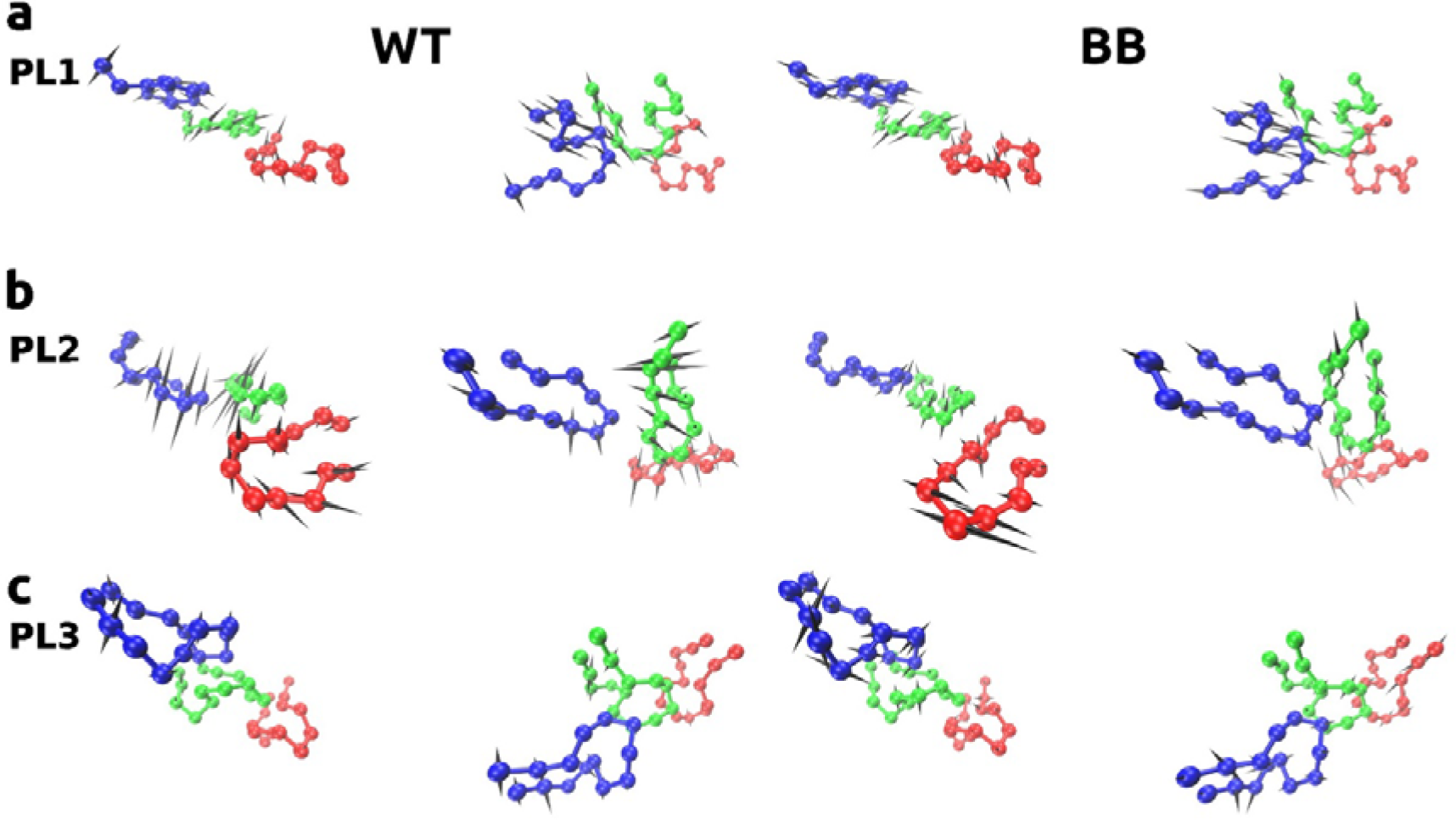
Effect of BB mutations pore loop dynamics. Motions associated with the PC1 eigenvector are shown for (a) PL1; (b) PL2 and (c) PL3 in protomers 2-4 (blue, green and red, respectively) of the wild-type ClpB (left panels) and of the BB variant (right panels). Top and side views are shown. Directions of motions are indicated using spikes. Motions of PL1 loops are less affected by mutations than those of the hexamer and PL2-3 loops. See also movies SM1-3.

To reveal allosteric paths connecting Walker B and pore loop sites within a ClpB subunit, we resort to a graph theory-based approach (39-47). The network comprises nodes representing individual residues, located at C_α_ positions, connected by edges whose lengths are weighted by the residue-residue cross-correlations (see Materials and Methods). It is important to note that the length of these path edges emphasizes the strength of allosteric coupling over the proximity in the Cartesian space of the residue pair. Thus, allosteric paths identified by this approach highlight the most effective propagation of allosteric coupling, therefore we focused here on the optimal path, with the shortest length, and suboptimal paths, with slightly longer lengths (see Materials and Methods). First, we determined up to 200 paths connecting the targeted mutation site from the Walker B regions of NBD1 (NBD1_B_), E279, or of NBD2 (NBD2_B_), E678, to the labeled residue of pore loop PL1 (A244), PL2 (A289) or PL3 (Y656). We examined three protomers from the high-resolution structure (9), and found that intra-protomer networks and allosteric paths are specific for each protomer. As shown in Tables S3 and S4, optimal paths identified in protomers of the wild-type ClpB are only slightly perturbed by the BB mutations. The limited changes to these paths are consistent with findings of previous studies, which noted that optimal paths are robust against perturbation (46). To obtain a broader understanding of the response of the allosteric network to these perturbations, and since current literature does not support the occurrence of a single allosteric pathway (46), we therefore studied also the suboptimal paths. As shown in Figures 4 and S5, the path length distributions of suboptimal paths have distinct behavior for the three pore loops. The PL1 distribution corresponds to stable allosteric coupling, with the similarity of path length distributions of wild-type and BB variants, quantified by the average overlap coefficient (OC) ∼ 0.44 ± 0.22 (see Fig. S5 j and Materials and Methods).

**Figure 4.**
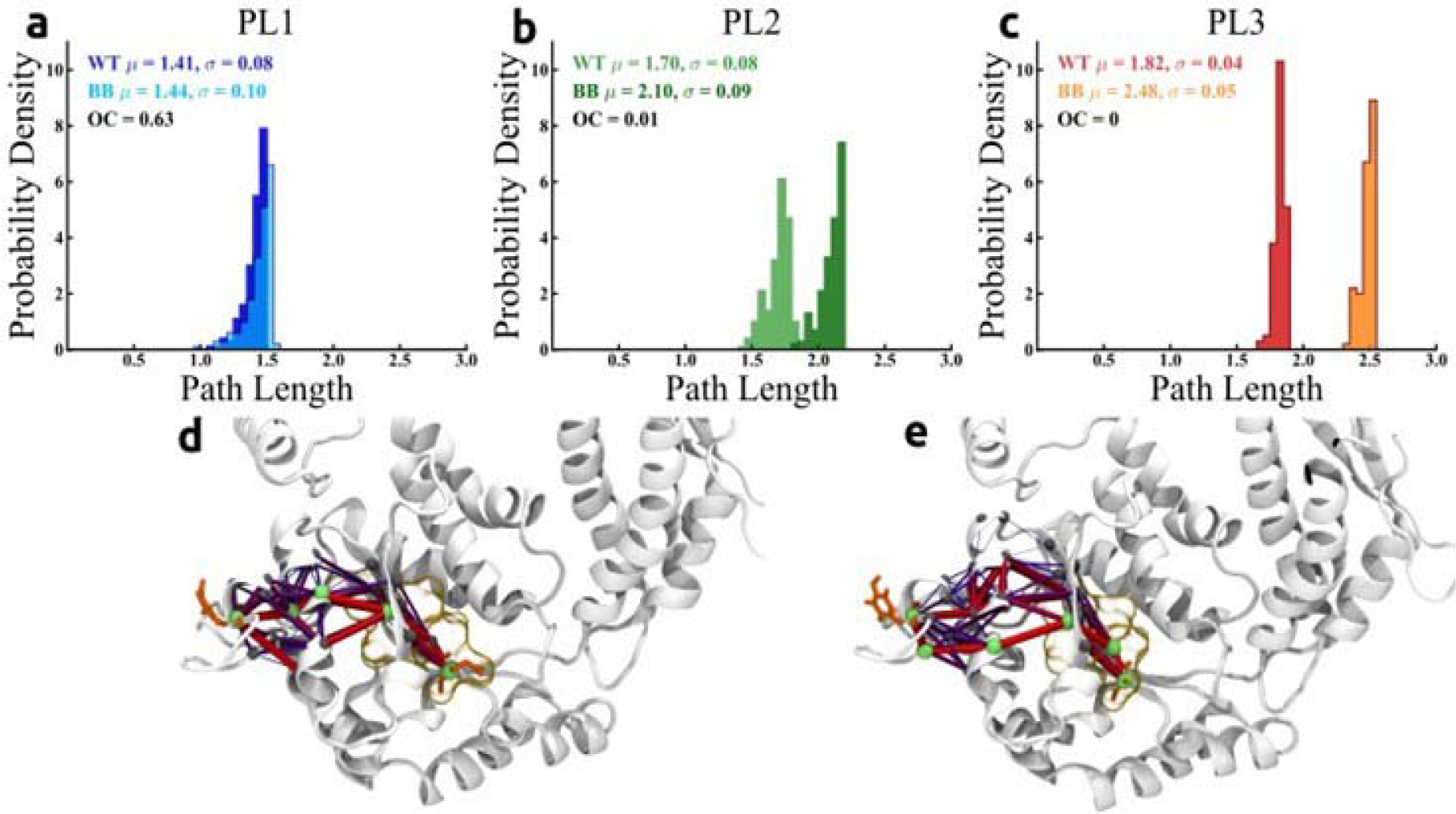
Computed allosteric paths connecting the Walker B and pore loop regions in ClpB. Probability density distributions of the 200 shortest paths between Walker B residues targeted by mutations, E279 in NBD1 and E678 in NBD2, and the pore loop residue labeled in FRET experiments, A244 in PL1, A289 in PL2, and Y656 in PL3 in one protomer of wt ClpB (*E. coli)* and BB mutant for **(a)** PL1 **(b)** PL2, and **(c)** PL3 wt and BB. The effect of perturbation on allosteric communication is weak in PL1, but strong in PL2 and PL3. Structural details of optimal and suboptimal paths are shown for PL3 **(d)** wt and **(e)** BB. Optimal paths (see Tables S3, S4), which have the shortest length, illustrate the strongest set of allosteric couplings between nodes (green) of the allosteric network, which mediate the signaling between the Walker B site and the pore loop. The optimal path is slightly perturbed by BB mutations whereas the ensemble of suboptimal paths (purple), which have longer path lengths, is strongly perturbed by mutations. Line thickness is proportional to the strength of the coupling.

By contrast, the PL2 distributions indicate a strong perturbation in the allosteric coupling to the NBD1_B_ region, with OC ∼ 0.16 ± 0.13 (Fig. S5 j). In the PL3 case, we found the largest shift in the strength of allosteric coupling compared with PL1 or PL2, with OC ∼ 0.09 ± 0.15 (Fig. S5 j). Thus, we find that BB mutants significantly altered the allosteric coupling of PL2 and PL3 dynamics, compared to PL1. The structural maps of allosteric paths (Fig. 4 and S6) illustrate the strong coupling of PL regions to active sites within the same NBD. The pattern of suboptimal paths that connect the Walker B region to PL1 remains largely unchanged in the BB mutant compared with the wild type (Fig. S6 a,b), whereas in the case of PL2 (Fig. S6 c,d) and PL3 (Fig. S6 e,f) it is strongly altered. Overall, our computational analysis supports the experimental results, which indicate a weaker effect of the BB mutations on PL1, and a stronger effect on PL2 and PL3.

### Absence of ATP hydrolysis strengthens κ-casein binding to pore loop 2

It was previously found that double-Walker B mutant of *E. coli* ClpB makes a stable interaction with model substrate-proteins, for which it was referred to as a “substrate trap” (48). As noted above, in our smFRET experiments, we see a concentration-dependent κ-casein-induced shift to state 1 (low-FRET), which indicates an increased population of state 1. We presume that at intermediate κ-casein concentrations, there exist both bound and unbound ClpB. However, because thousands of such molecules are analyzed in the measurement, we obtain an effective concentration-dependent shift (Fig. S11), and can make use of this shift to estimate and compare κ-casein binding affinity to each pore loop by conducting smFRET κ-casein titrations.

To this end, we have conducted smFRET measurements for PL1, PL2 and PL3 constructs as well as for their BB mutants in the presence of increasing concentrations of unlabeled κ-casein, and carried out H^2^MM analysis of these datasets with two fixed states, as described before. We have calculated the ratios of H^2^MM-derived populations of state 1 (at low-FRET) with and without casein. Plots of these ratios against the concentrations of κ-casein yielded saturation curves for all three types of pore loops (Fig. 5). Fits of these curves to a simple binding isotherm revealed micromolar dissociation constants for PL1 and PL3 (2.9 μM and 0.9 μM, respectively, Table S5). In this analysis, neither PL1 nor PL3 showed dramatic differences in κ-casein binding between the non-mutated wt and the ATP-hydrolysis deficient BB constructs. In contrast, however, there was a surprising 40-fold increase in the binding affinity for PL2 upon abolishing ATP hydrolysis, and the derived K_d_ changed from 11.3 μM in PL2 wt to 0.3 μM in PL2 BB construct. Based on this difference, it is plausible that the increased substrate binding to PL2 is involved in the ‘substrate trap’ effect noted in the past (48). In our data, in the high-affinity PL2 BB construct, the high-FRET state (state 2), corresponding to the ‘down’ conformation of PL2, becomes more populated than in PL2 wt (0.67 ± 0.02 vs. 0.56 ± 0.01 in the wt, Table S6). Possibly, this conformation allows maximizing contacts with the substrate-protein and this way stabilizes its binding, and acts to prevent its back-slipping.

**Figure 5.**
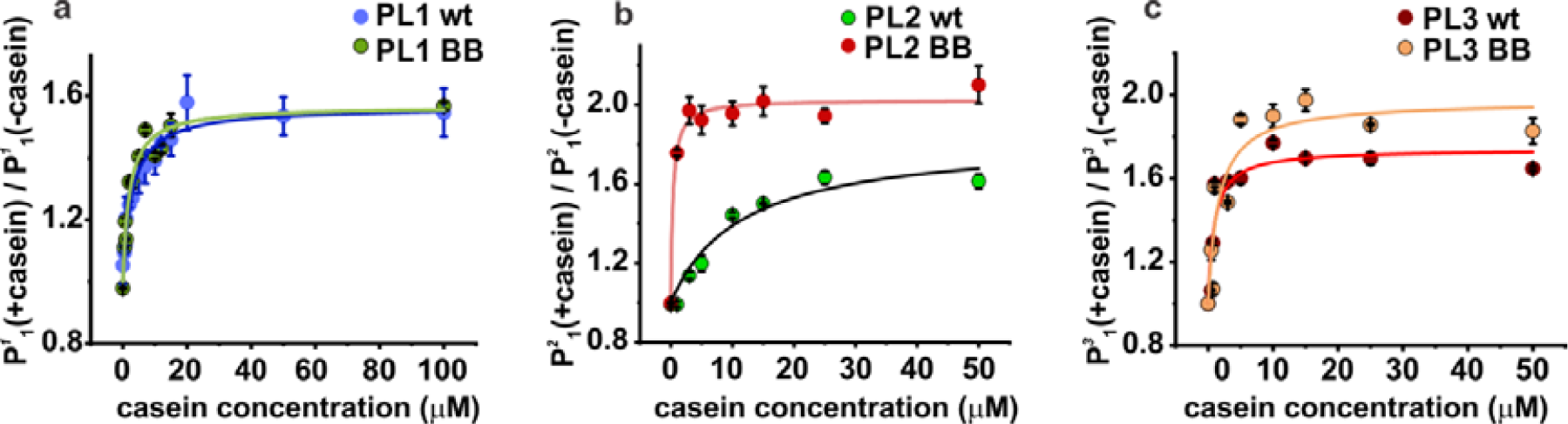
smFRET κ-casein titration experiments with wt and BB constructs. The average ratio of state 1 population with κ-casein to that without κ-casein is plotted (as circles) against casein concentration. Error is SD (n=3) for PL1 and PL3 datasets, and SD (n=2) for PL2 datasets. Solid lines are fits to a binding model (details in Table S5).

### Both NBD1 and NBD2 impact pore-loop dynamics

In light of our results with the BB mutants, we set to determine whether and how hydrolysis in NBD1 and NBD2 separately affects pore-loop dynamics. In particular, we wanted to find out if the nucleotide state of each NBD can affect only the pore loops that are located within that NBD, or if this effect is longer-ranged.

To study the coupling of NBD1 and NBD2 to pore loops individually, we introduced single mutations into Walker B or Walker A motifs of either NBD. We generated single-NBD mutants which could not hydrolyze ATP in NBD1 (with mutation E271A, denoted as “B1”), or in NBD2 (E668A, “B2”). We also prepared mutants with abolished ATP binding to NBD1 (K204T, “A1”) or to NBD2 (K601A, “A2”). As a positive control for ATPase activity, we prepared a hyperactive mutant, which is expected to show elevated ATP hydrolysis rate due to an effect on its middle domain (K347A, “hyper”) (49). These constructs were homogeneously assembled, and their ATPase activity was found to be altered relative to the wt as expected, either lowered in the Walker mutants or increased in the hyperactive construct (Fig. S12). Having checked their bulk properties, we analyzed these constructs by smFRET spectroscopy.

All constructs displayed broad FRET efficiency histograms (Fig. 6, S13, S14) with a shift to low FRET efficiency values upon κ-casein addition, and fast transition rates from the H^2^MM analysis, on the same timescale as for the non-mutated pore-loop constructs (Tables S6). However, the populations of the two states were affected, as indicated by the shifts in the FRET efficiency histograms as well as by the H^2^MM-derived effective equilibrium coefficients, *K*_*i*_s, as is summarized in Figure 6 a-c.

**Figure 6.**
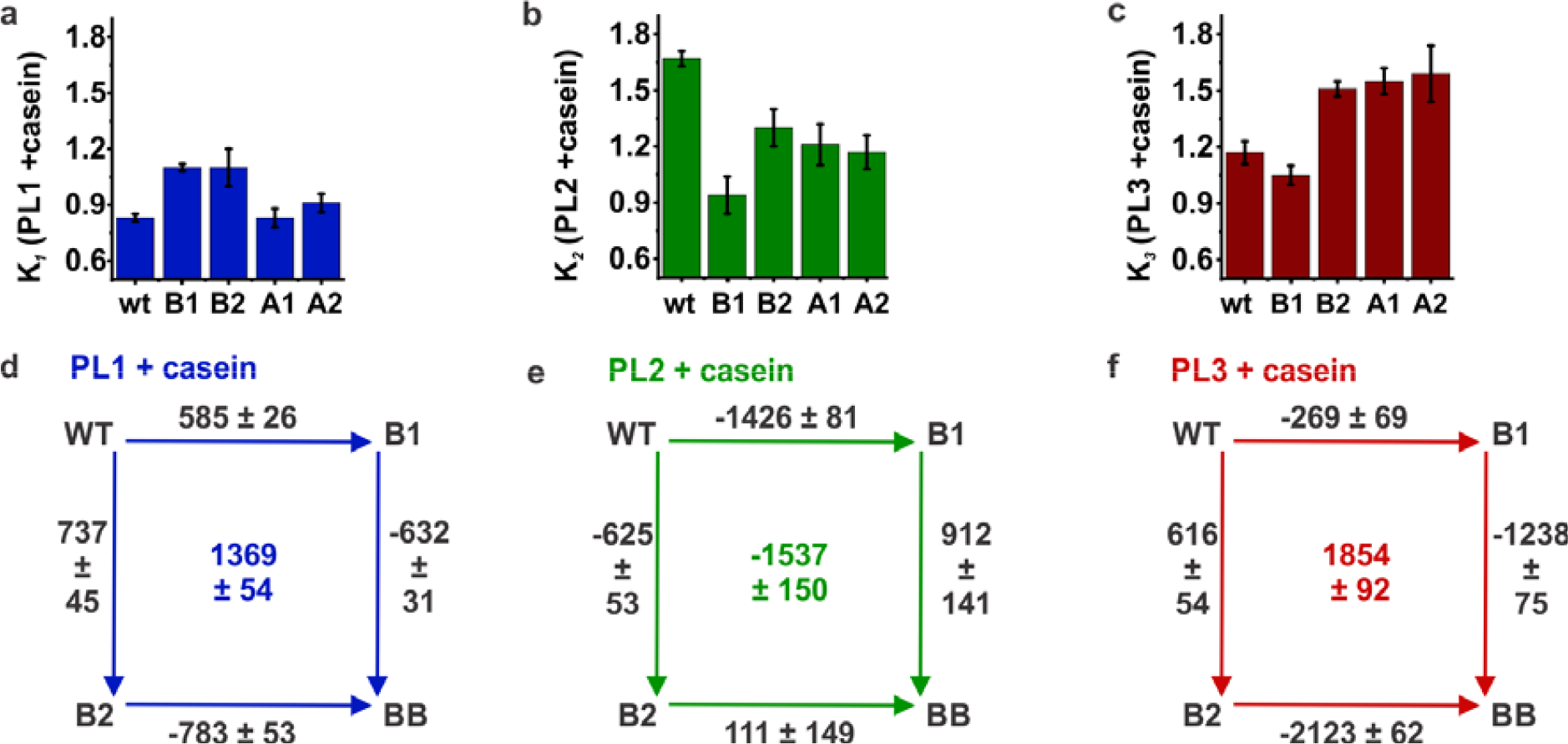
Effect of single NBD mutations on PL1, PL2 and PL3. **(a-c)** H^2^MM-derived equilibrium coefficients, s, for sets of single Walker A/B mutants of PL1 **(a)**, PL2 **(b)** and PL3 **(c)**, all measured with 25 μM κ-casein (average values, errors are SD, n=2-3 repeats). See Table S6 for values without κ-casein and Figures S13-S14 for complete sets of smFRET histograms with single NBD mutants. **(d-f)** Double-mutant cycles for PL1, PL2 and PL3 with κ-casein (25 μM). The values along the edges are the average changes in free energy upon mutation, (in J.mol^-1^), calculated from the H^2^MM-derived s detailed in Methods. The values at the centers of the squares are the average coupling energies, defined as the differences between free energy changes of opposing edges (see Materials and Methods). Error values are from the propagation of the standard errors in. The cycles without κ-casein are in Fig. S15.

PL1 was least affected by the mutations in the NBDs, and both B1 and B2 mutations slightly increased the population of the low-FRET state (‘up’ conformation, Figs. 6, S13, S14), even though this effect was surprisingly lost in the BB mutant (Fig. S1). The observation that this pore loop is affected by hydrolysis in either NBD indicates the presence of long-range allosteric communication in ClpB. PL2 was affected by all mutations in the same way: they all led to the increase in the population of the high-FRET (‘downward’) conformation which might aid with the substrate-protein binding. For example, abolishing ATP hydrolysis in NBD1 or in NBD2 caused *K*_2_ to change from 0.79 ± 0.02 to 0.37 ± 0.01 (in PL2 B1 construct) or to 0.41 ± 0.01 (in PL2 B2), respectively (Tables S6). Thus, the effect is more prominent when ATP is bound and not hydrolyzing in NBD1 than in NBD2, suggesting that there is stronger coupling of PL2 to NBD1 than NBD2. Nevertheless, the dynamic equilibrium of PL2 can be regulated by the nucleotide state of either NBD.

In PL3, the dynamics were strongly affected by the studied mutations, suggesting a significant mechano-chemical coupling of this pore loop to the NBDs. As already mentioned, in the absence of ATP hydrolysis in both NBDs (BB construct), the population of high-FRET (‘up’) state was increased (Fig. 2). In contrast, B2, A2 and A1 mutations led to increase in the low-FRET state (‘downward’ conformation relative to wt), while B1 mutation had almost no effect (Fig. 6 and Figs. S13, S14). In the B2 construct, changed from 0.50 ± 0.02 in wt to 0.79 ± 0.03 in B2 mutant without κ-casein, and the difference was also present with κ-casein addition (1.17 ± 0.06 in wt and 1.51 ± 0.04 in the mutant, Tables S6). According to these results, the dynamic equilibrium of PL3 can be shifted to either its up or down conformation depending on the nucleotide state of the NBDs. In the ATP-bound state in the absence of hydrolysis, PL3 favors a strained upward conformation. Once ATP hydrolysis occurs in the upper NBD (B2 mutant), or nucleotide gets released from either the upper or the lower NBD (A1 and A2 mutants), this stabilization is lost and the low-FRET (‘downward’) state becomes dominant. This population redistribution in PL3 is likely to be crucially involved in the substrate-protein translocation by ClpB.

### Double-mutant cycles expose significant coupling between NBDs

To further analyze the contributions of Walker B mutations in either NBD or in both, we employed double-mutant cycles (DMCs) (37). We calculated the free energy differences for each construct, Δ*G*_*i*_, by using the H^2^MM-derived *K*_*i*_ values. We subsequently calculated the free energy change upon each mutation, ΔΔ*G*_*i*_, as the difference between the two corresponding Δ*G*_*i*_ values (detailed in Materials and Methods) and plotted thermodynamic cycles where the resulting ΔΔ*G*_*i*_ values are included along the edges (Fig. 6d-f, Fig. S15). In these DMCs, the values along the opposite edges are not equal in all cases, indicating thermodynamic coupling (37). Considering that both the ATP-binding sites and the pore loops are separated in space and thus cannot interact directly, this result indicates long-range coupling. Interestingly, the coupling energies, calculated as the differences in ΔΔ*G*_*i*_ between opposite edges (see Materials and Methods), are positive for PL1 and PL3, both with and without κ-casein, suggesting that the introduction of one Walker mutation has a stabilizing effect on the other Walker mutation. The interaction energy for PL2, though, is negative.

## Discussion

The details of the translocation mechanism by AAA+ proteins are incompletely understood to date. Multiple experimental findings, as mentioned in the Introduction, indicate that the functions of these molecular machines might involve complex conformational changes on multiple length- and time-scales. Apart from the slow ATP-dependent sequential movements of protomers, as inferred from cryo-EM images, intra-protomer motions, such as fluctuations of pore loops, can also significantly contribute to the function of these proteins.

In this study, we selectively monitor the intra-protomer motions of individual pore loops along the axial channel of ClpB and how they are affected by nucleotide-binding site mutations. Consistent with our recently published experimental results (28) and with theoretical predictions (36), we find that the pore loops are moving on the microsecond timescale. Furthermore, we see that the ratio of the ‘up’ and ‘down’ conformations of the pore loops along the axial channel is modulated in response to the changes in the ATPase state of each of the NBDs, indicating that the pore loops and the NBDs are allosterically coupled. Interestingly, suppression of ATP hydrolysis in both NBDs leads to a perturbation of PL2 and PL3, but not PL1, suggesting a different role for the latter. To characterize these couplings, we perform molecular dynamics simulations of ClpB, and calculate the allosteric paths that connect the NBDs to the pore loops. These simulations show clear perturbations of the allosteric network and of the pore loop dynamics due to ATP-hydrolysis abolishing mutations, in good agreement with our experimental findings. Surprisingly, each of the NBDs can affect all pore loops, even if they do not belong to the same NBD, suggesting a long-range allosteric regulation in ClpB. Indeed, by using our experimental equilibrium coefficients in a double-mutant cycle analysis, where the cycles include wt, BB and the single B1 and B2 mutants, with the mutations being present in all six protomers of the hexamer, we find that the effects of the single mutations are not independent, with a significant energy of interaction coupling them (Figs. 6, S15) (37). Allosteric communication between the two nucleotide binding sites, NBD1 and NBD2, in ClpB (50-52) and related proteins (53) could previously be inferred from functional studies of ATP hydrolysis. What we report here is, in contrast, an uncharacterized form of such a coupling, operating through the long-range effect on the pore loops and leading to a direct effect on the mechanism of protein translocation.

Our experiments demonstrate quite remarkably that the conformational dynamics of PL3 show differential response to mutations in either of the two NBDs (results are summarized in Fig. 7). This pore loop is highly conserved within the AAA+ family (5), suggesting that a similar NBD-pore-loop communication might exist in other AAA+ members. It is likely that the ATP-dependent change in the conformational equilibrium of PL3 facilitates the translocation of substrate-proteins across the central pore of ClpB. These results are in agreement with the previously reported evidence for nucleotide-dependent pore-loop motions in ClpB (21) and ClpX (25,26), and are consistent with the captured large-scale (17 Å) longitudinal motions of the pore loops in a DNA-unwinding AAA+ hexamer (12). Thus, the up/down motions of the substrate-binding pore loops might be conserved in the AAA+ machines and possibly serve to pull the substrates across the hexameric rings. Our analysis shows that these up/down motions occur on the microsecond timescale but can be affected by the changes in the nucleotide state of the protein, which is modulated on the timescale of ATP binding and hydrolysis. Altered microsecond pore loop motions, as well as their effects on average populations of the up and down states of the pore loops, may ultimately affect the dynamics of substrate translocation.

**Figure 7.**
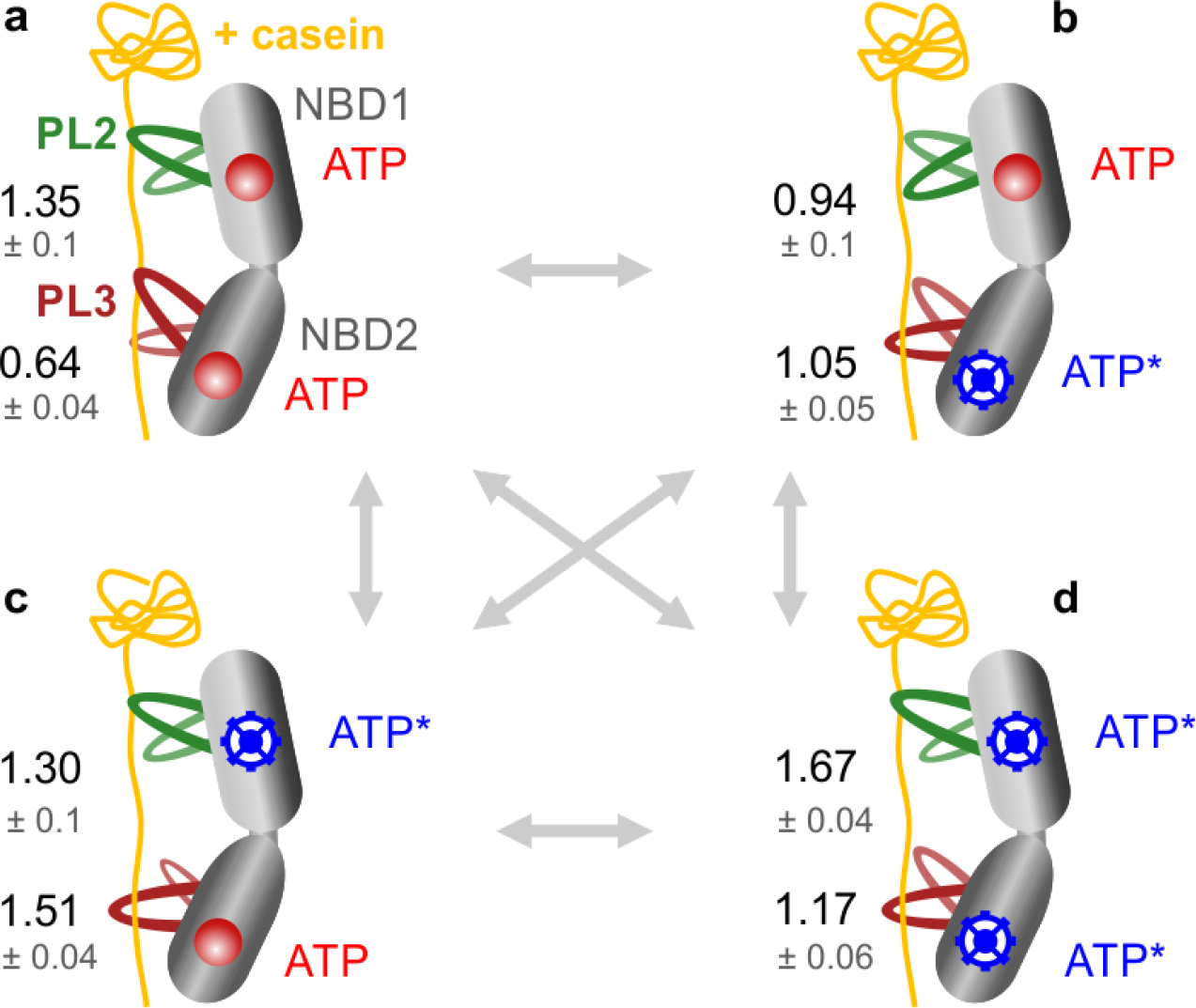
smFRET results suggest an ATP-dependent modulation of the pore-loop dynamics. **(a-d)** States of ClpB monomer are schematically shown, with bound κ-casein (in yellow), and pore-loops PL2 and PL3 in green and dark-red, respectively. PL1s are omitted for simplicity but included in Fig. S16. The ATPase states of the NBDs are depicted as follows: red circles are bound ATP molecules not undergoing hydrolysis (ATP arrested state), blue wheels – bound ATP undergoing hydrolysis, which corresponds to a mixture of ATP/ADP, although ATP is in excess in these measurements (2 mM). As a consequence of altered microsecond rates, PLs favor either an up or a down conformation depending on the ATPase state of the NBDs, and the size of the PLs on the scheme reflects their state occupancy from the H^2^MM analysis. Numbers are the average H^2^MM-derived equilibrium coefficients, *K*_*i*_ (errors are SD, n=2-3). The conformational equilibrium of the pore loops periodically switches between the up/down states in response to the changing ATPase state of the machine, and this might facilitate substrate-protein translocation.

Thus, the coupling of pore loops to NBDs involves two timescales and is consistent with the Brownian-ratchet mechanism (54,55) for translocation by ClpB, as has been proposed in our preceding work (28,56). In this mechanism, the change in the population ratio of the pore-loops that occurs upon ATP binding/hydrolysis corresponds to a shift between two free-energy surfaces, flat and structured, and results in a unidirectional, rectified substrate-protein translocation (55). Because of these two distinct timescales characterized, we can infer that the translocation by ClpB is likely to be faster than what can be expected based only on its ATP hydrolysis rate, in agreement with the recent study by optical tweezers (19). Indeed, since pore-loop motion is not fully coupled to substrate translocation, the microsecond motion of the former can readily lead to millisecond substrate translocation, as reported in that paper.

Our approach allows us to experimentally measure the substrate-binding affinities of distinct pore loop types in ClpB. We find that pore loops 1 and 3 and their ATP-hydrolysis deficient BB mutants bind to the substrate-protein κ-casein with similar affinities. In contrast, the ATP-hydrolysis deficient PL2 BB mutant binds to the substrate ∼40 times stronger than the wild type PL2. Our ability to measure substrate-protein affinities to distinct pore loop types is entirely novel and lends support to previous findings that double Walker-B mutations result in stronger substrate-protein binding to ClpB (48), and might offer a more nuanced explanation of these bulk observations. Considering that PL2 is the least conserved pore loop, its ATP-hydrolysis-dependent change in protein-binding affinity might act as a very effective filter for substrate-protein selection before they get passed to the lower NBD2 and bind to PL3. The latter is conserved in the AAA+ family (5), which might imply that substrate-protein recognition by PL3 is not highly specific. Indeed, ClpX, which contains a tyrosine pore loop analogous to PL3 of ClpB, was previously demonstrated to translocate a range of polypeptide substrates independently of the steric or chemical properties of their side chains (57). It might be that if the bound substrate-protein passes the ‘filtration’ by the less conserved PL2 within NBD1, it subsequently gets transferred into NBD2 and is committed to complete translocation across the axial pore of ClpB. Similarly, selectivity during substrate-protein recognition in ClpX is ensured by its RKH and pore-2 loops, which are poorly-conserved in other AAA+ machines (25,58).

By summarizing our data in Figure 7, we can speculate on how the pore loops in ClpB facilitate the ATP-dependent substrate-protein translocation. Given that our experiments are performed under saturating ATP concentrations (2 mM), that ATP binding is stochastic and that ATP hydrolysis is orders of magnitude slower than pore-loops motions, all four states of ClpB that are depicted in Figure 7 are likely to be present. From our results, it is plausible that ATP binding and hydrolysis events drive substrate-protein translocation by allosterically modulating the dynamic equilibrium of PL2 and PL3.

Although we have not directly measured substrate-protein translocation by ClpB, by considering the results from both our preceding (28) and current study, we can make assumptions on how ATP coordinates substrate-protein translocation. In particular, we can suggest that the changes in the dynamic equilibrium of PL3 are directly coupled to ATP-dependent protein translocation. Previously, we established that PL3 dynamics are correlated to bulk disaggregation activity of ClpB (28). Furthermore, we observed both previously (28) and currently that PL2 is ATP-dependent but does not correlate to any bulk activities, neither disaggregation nor ATPase. We therefore assume that PL2 functions as a pawl, which acts to ensure unidirectional substrate-protein translocation. Regarding PL1, we previously found it to be nucleotide-type independent, and do not observe effect on its dynamics upon abolishing ATP hydrolysis (in the BB construct). These results are consistent with the recent results from molecular dynamics simulations (36). Although the dynamics of PL1 are independent of the machine’s nucleotide state, we still expect this pore-loop type to be important for translocation, since it is likely to be essential for the initial substrate binding. Indeed, we also previously found that PL1 dynamics correlate with bulk disaggregation activity (28), and therefore expect PL1 to be crucial for the substrate-protein engagement and translocation, though we do not include it in Fig. 7 for simplicity. We include a summary of our results for all three PLs in Supplemental Information (Fig. S16) along with a further discussion of the potential translocation mechanism (Fig. S17).

Using Walker B mutants, we now find that PL2’s affinity for substrate-protein is dramatically increased when ATP is bound to both NBDs but not undergoing hydrolysis (BB construct or ATP arrested state, Fig. 7a). This result indicates that simultaneous ATP binding to both NBDs, before ATP hydrolysis takes place, likely acts as a high-affinity state of ClpB that ensures substrate-protein binding and possibly prevents it from back-slipping.

At the same time, in this ATP arrested state (Fig. 7a), PL3 is stabilized in an upward conformation, which might aid with the substrate-protein transfer into NBD2. PL3 binds the substrate-protein, either to prepare for pulling or to act as pawl. Either upon ATP hydrolysis in NBD2 (Fig. 7b) or in both NBDs (Fig. 7d), where PL3 samples the up/down states almost equally, it acts by pulling; whereas upon ATP hydrolysis in NBD1 (Fig. 7c), where is favors the downward state, it likely acts as a pawl. Alternatively, the fast up and down transitions of PL3 (in Fig. 7b and d) perhaps act to allow PL3 to reconfigure its binding to the substrate (this is similar to our findings with the enzyme, which opens and closes multiple times to optimize substrate configuration (31)). ATP hydrolysis in NBD1 (Fig. 7c) stabilizes the down conformation of PL3, likely promoting substrate pulling toward the exit of the channel. To note, the stabilization of the downward conformation, seen upon ATP hydrolysis in NBD1 (Fig. 7c), also occurs upon ATP release from either NBD (Fig. S16).

ATP hydrolysis in NBD2, while ATP is bound to NBD1 (Fig. 7b), allosterically modulates PL2, slightly stabilizing its downward conformation. This action likely allows PL3 to pull or reconfigure while PL2 acts as a pawl. As mentioned above, ATP binding to NBD2, while NBD1 is undergoing ATP hydrolysis (Fig. 7c), allosterically modulates PL3, stabilizing its downward conformation. This action likely allows PL2 to release its contact form the substrate and allows PL3 to act as a pawl or to pull on the substrate-protein.

## Conclusion

To conclude, our results establish a considerably long-ranged allosteric communication between the ATP binding sites of ClpB and its pore loops, which are responsible for substrate translocation, thus establishing the pore loops as active participants during ATP-dependent translocation. Importantly, while previous studies have indicated that pore loops bind to substrate proteins and possibly undergo conformational changes, this is the first direct and detailed biophysical characterization of the ATP-dependent regulation of the real-time dynamics of three distinct types of pore loops by each of the two ATP binding sites. Remarkably, we find a significant interaction between the two ATP-binding sites in their effect on pore-loops. This coupling modulates the dynamic equilibrium between the rapidly exchanging up/down states of the pore loops, thereby facilitating the ATP-dependent unidirectional translocation of protein substrates. Our smFRET measurements, coupled with molecular dynamics simulations and mutational analysis, allow us to obtain a comprehensive picture of these allosteric interactions and to propose how dynamics on the slow time scale of ATP hydrolysis affects the much faster motions of protein-translocating structural elements. These results, and the methods presented here, should be relevant to the studies of multiple additional AAA+ proteins.

## Supporting information

Supplemental Information (revised)

## Author Contributions

M. I. designed and performed research, analyzed data and wrote the paper. H. M. designed and performed research and contributed analytic tools. A. D., Z. Z. and G. S. designed research, contributed analytic tools and analyzed data. I. R. and G. H. designed research and wrote the paper.

## Acknowledgements

We thank Prof. Amnon Horovitz for the advice on the DMC analysis. M.I. was the recipient of an EMBO Long-Term Fellowship (ALTF 317–2018) and the IASH Fellowship for International Postdoctoral Fellows. H.M. was supported by Planning & Budgeting Committee of the Council of Higher Education of Israel. G.H. was funded by the European Research Council (ERC) under the European Union’s Horizon 2020 research and innovation programme (grant agreement No 742637), by NSF-BSF grant No 2021700 and by the Grand Center for Sensors and Security. G.H. is the incumbent of the Hilda Pomeraniec Memorial Professorial Chair. G.S. was funded by NSF-BSF grant MCB-2136816. This work used Bridges-2 resources at the Pittsburgh Supercomputer Center through allocation MCB170020 to G.S. from the Advanced Cyberinfrastructure Coordination Ecosystem: Services & Support (ACCESS) program, which is supported by National Science Foundation grants #2138259, #2138286, #2138307, #2137603, and #2138296.

## Data Availability

Data generated in this work are available upon reasonable request.

## Supplemental Information

Supplemental smFRET Data Analysis Details, Supplemental Computational Methods, Supplemental Figures S1-S17, Supplemental Movies SM1-3 and Supplemental Data Summary Tables S1-S6 are available online.

