## Supplemental Information (revised) for "Single-molecule FRET probes allosteric effects on protein-translocating pore loops of a AAA+ machine"

### Single-molecule FRET probes the allosteric effect of ATP on the protein-translocating pore loops of a AAA+ machine

Movie SM3. 24
Supplemental Data Summary Tables: 25
Table S1. FRET efficiency values of the two pore loop states as obtained from the global $H^2MM$ analysis of wild-type pore-loop constructs, measured with or without κ-casein 25
Table S2. Cumulative overlap of the top 2 principal components of pore loops in the BB variant over the top 10 principal components of the corresponding pore loops in wild-type ClpB. 25
Table S3: Optimal paths between Walker B sites and pore loops in NBD1, PL1 and PL2, in wild type ClpB and BB mutants, derived from the MD simulations of *E. coli* ClpB. 26
Table S4: Optimal paths between Walker B sites and pore loops in NBD2, PL3, in wild type ClpB and BB mutants, derived from the MD simulations of *E. coli* ClpB. 26
Table S5. Substrate κ-casein binding constants to individual pore-loop mutants, derived from $H^2MM$ analysis of single-molecule κ-casein titration measurements. 26
Tables S6: $H^2MM$ Analysis Outputs. 27
PL1: WT and double mutants. 27
PL2: WT and double mutants. 27
PL3: WT and double mutants. 27
PL1: Single-NBD mutants. 28
PL1: Other conditions. 28
PL2: Single-NBD mutants. 29
PL2: Other conditions. 29
PL3: Single-NBD mutants. 30
PL3: Other conditions. 30
Supplemental References 31

**Supplemental smFRET Data Analysis Details:**

#### Fixed states for $H^2MM$ analysis

As in our previous paper (1), here we characterized pore-loop dynamics using the $H^2MM$ algorithm, a photon-by-photon hidden Markov modeling approach capable of quantifying microsecond-timescale dynamics in single-molecule FRET data (2). We previously determined two discrete FRET efficiency states to be essential and sufficient for the modeling of pore-loop dynamics. This was done by first using a model with 9-10 equally spaced FRET efficiency states, and finding that the derived free energy profiles showed only two major minima, which we assigned to 'up' and 'down' conformations of the pore loops (1). Therefore, here we also used the two-state $H^2MM$ model for all data analysis presented in this study. Furthermore, we previously globally analyzed wild-type pore-loop constructs measured with and without a saturating concentration of κ-casein (20 µM), assuming that the two FRET efficiency states for each pore loop remain the same under both conditions (1). In the present work, the FRET efficiency values for each pore loop from this global analysis (listed in Table S1) are used as fixed inputs for the modelling of the datasets of all generated mutants of the corresponding pore-loop construct. This is done in order to simplify our analysis and to ensure the most direct comparison between the state population redistributions as a consequence of introduced mutations.

#### $H^2MM$ analysis validations

This $H^2MM$ analysis was validated through recoloring and segmentation analyses, which were described in detail in our preceding study (Fig. S5 in Mazal *et al*. (1)). Briefly, in the statistical recoloring analysis, described previously (1,3), the arrival times of photons, detected in the smFRET experiment, are kept unchanged, whereas the identity ('color') of photons, i.e., whether they result from the emission by AF488 or AF595 dye, is erased. The photons are subsequently 'recolored' using a Monte-Carlo simulation based on the model parameters derived from the $H^2MM$. Average FRET efficiencies for every burst are calculated, and a histogram is generated and compared to the experimentally measured histogram. Agreement between the two histograms indicates valid $H^2MM$ modeling (Fig. S2). For the segmentation analysis, previously described (1), the most probable sequence of states in each burst is used, as obtained from the $H^2MM$ analysis and the Viterbi algorithm (which is an algorithm that delineates the single best state sequence in each trajectory). Based on the assignment of each segment within the burst to state 1 or state 2, FRET efficiency histograms are calculated. It is expected that the calculated histograms will show a clear separation into two states for a valid two-state model (Fig. S2). We further validated the two-state analysis by calculating dwell time distributions, following a previously described procedure using a custom-written likelihood-weighted segmentation algorithm (3). This procedure analyzes the distribution of dwell times that are spent by the pore loops at each of their two states. For any photon trajectory, this analysis does not only take into account the most probable sequence of states (classically obtained by the Viterbi algorithm). Rather, every possible sequence of states contributes to the dwell time a fraction of a count that is equal to the likelihood of this sequence. Integrated dwell time distributions are computed and fitted to single exponential functions to derive the transition rates. Agreement between transition rates, derived from $H^2MM$, with the rates from the dwell time analysis provides the strongest indication for valid modeling (Tables S6).

**κ-casein addition does not impact the photophysical properties of Alexa Fluor dyes**

In our previous smFRET studies of ClpB, we verified that κ-casein addition has no effect on the photophysical properties of Alexa Fluor dyes. We reported that the presence of 25 μM of unlabeled κ-casein in our measurements has no effect on the mean numbers of photons per burst, mean photon rates or average burst lengths (as detailed before (4)). Furthermore, we observed identical FRET efficiency histograms in the presence and in the absence of κ-casein for two separate ClpB constructs with Alexa Fluor dyes located on rigid positions within NBD1 and NBD2 (1,4).

#### Supplemental Computational Methods:

##### Molecular dynamics simulations

Wild-type ClpB simulations involve solvation of the ATPase in a cubic box with dimensions ~ 170 × 170 × 170 $Å^3$, with 138913 water molecules represented using the single point charge (SPC) model and neutralized by adding 105 Na ions. The simulation box includes a total of 452304 atoms. Periodic boundary conditions (PBC) were applied in the three dimensions. For ClpB mutants, the protein structure was solvated in a dodecahedral box with dimensions ∼ 115 × 163 × 163 $Å^3$ with 85780 water molecules, represented using the single point charge (SPC) model. 93 Na ions were added to the system to neutralize charges, resulting in a system with 292845 atoms. Energy minimization of the solvated systems was performed using the steepest descent algorithm for 50000 steps with convergence achieved when the maximum force reached a value smaller than 1000 kJ/ (mol · nm). Next, two equilibration steps were performed. First, NVT simulations were performed for 500 ps using the leapfrog integrator, with T = 300 K, and harmonic restraints applied to heavy atoms of the protein with a spring constant of 1000 kJ/ (mol · $nm^2$). In the second equilibration step, NPT simulations of the restrained system were performed for 500 ps with the pressure maintained at the constant value of 1 atm using the Parrinello Rahman algorithm (5). The time step in all molecular dynamics (MD) simulations was 2 fs. After removal of restraints, five unbiased NPT simulations of trajectories were performed for 150 ns (wild-type) and 150 ns (BB mutant). Thus, wild-type simulations, including previous data (6), comprised 150 ns for each MD trajectory. For analysis purposes, the first 10 ns of each trajectory were not included, and data frames were saved every 100 ps. Equilibration of simulation trajectories is assessed using root-mean-square deviations (Fig. S7). The stability of nucleotide location is evaluated by determining the distance between the center of mass of the nucleotide and the $C_\alpha$ atom of the Walker B mutation position within the same nucleotide binding site (Fig. S8), and the stability of nucleotide-ClpB interactions is quantified by calculating the total interaction energy, comprising Coulomb and Lennard-Jones terms for atom pairs within the cutoff distance of 12 Å, between each nucleotide and the ClpB hexamer (Fig. S9).

##### Dynamic Cross-Correlation Matrix

We used the Bio3D package (7) to determine the Dynamic Cross Correlation Matrix (DCCM) of position fluctuations of $C_\alpha$ atoms of protein residues, which quantifies the time-dependent residue-residue directional correlations. DCCM is an N × N matrix, where N is the number of residues, where each element $C_{ij}$ corresponds to the dynamic cross-correlation between residues i and j:

$$C_{ij}(t) = \langle \triangle \mathbf{r}_i(t')\cdot\triangle \mathbf{r}_j(t')\rangle / (\langle\|\triangle \mathbf{r}_i(t')\|^2\rangle\langle\|\triangle \mathbf{r}_j(t')\|^2\rangle)^{1/2}$$

Here, $\triangle \mathbf{r}_i (t') = \mathbf{r}_i (t') - \langle \mathbf{r}_i \rangle$ denotes the instantaneous position fluctuation of residue i from its mean. Ensemble averages over all time frames up to time t and all trajectories are indicated by $\langle\cdot\rangle$. $C_{ij}$ values range from -1 to 1, with motions of i and j atoms in the same direction corresponding to positive $C_{ij}$ values, and motions in opposite directions to negative values. Convergence of the DCCM matrix was assessed using the mean square distance R(t) between correlation values of residue pairs at successive times, $R(t) = (1/N_p) \sum_{(ij)} (C_{ij}(t) - C_{ij}(t-\tau))^2$, where $N_p$ is the number of residue pairs and the time interval $\tau = 10$ ns(5). Here, $C_{ij}$ is evaluated using data frames up to the total simulation time per trajectory $t \leqslant 150$ ns. As shown in Figure S10, DCCM convergence for wild type and BB simulations is achieved within approximately 80 ns.

**Principal component analysis**

To capture important modes of pore loop dynamics, we performed principal component analysis (PCA) applied to the $C_\alpha$ positional fluctuations of the wild-type ClpB and the BB variant. In separate calculations, we performed PCA for each of the PL1, PL2, or PL3 pore loop types in protomers 2, 3 and 4. Protomers 2-4 are selected for the analysis as both NBDs are in an active (ATP-bound) state in the pre-hydrolysis ClpB structure (PDB-6OAX (8)). Although protomer 5 is also bound to ATP in both NBD1 and NBD2, it was not included in our analysis, since its greater flexibility, due to the weak lateral interface with protomer 6, is overemphasized in the truncated ClpB variant given the absence of the middle domains. Normalized eigenvectors (PCs) and their associated eigenvalues were obtained by diagonalization of the covariance matrices of position fluctuations $\langle \triangle\mathbf{r}_i \cdot \triangle\mathbf{r}_j \rangle$ using Python packages MDTraj (9) and NumPy (10). We compared the similarity of PC motions within the essential subspaces of the wild-type and BB variants by calculating the overlap of two eigenvectors, the cumulative overlap (CO) of one eigenvector of one subspace and eigenvectors of the other subspace and root-mean-square inner product (RMSIP) between eigenvectors of the two subspaces. The overlap between the directions of a wild-type PC and a BB PC is (11)

$$O_{ij} = |\ \mu_i^{wt} \cdot \upsilon_j^{BB}\ |$$

where $\mu_i^{wt}$ is the *i^th^* PC of the PL in wild-type ClpB, and $\nu_j^{BB}$ is the $j^{th}$ PC of the same PL in the BB variant. The cumulative overlap between the first *k* PCs of the PL in wild-type ClpB and the $j^{th}$ PC of the PL in the BB variant is defined as (12)

$$CO_j(k) = \left( \sum_{i=1}^{k} O_{ij}^2 \right)^{\frac{1}{2}}$$

The CO provides information about the extent to which a single BB PC is represented within the motions captured by the first *k* wild-type PCs. RMSIP measures the overlap between subspaces spanned by the first *m* PCs of wild-type and BB variants by using (12,13)

$$RMSIP(m) = \left( \frac{1}{m} \sum_{j=1}^{m} \sum_{i=1}^{k} O_{ij}^2 \right)^{1/2} = \left[ \frac{1}{m} \sum_{j=1}^{m} (CO_j(k)^2) \right]^{\frac{1}{2}}$$

**Optimal and suboptimal path analysis**

We performed the path analysis in protomers 2-4 (PDB-6OAX (8)) and determined the optimal and suboptimal intra-domain paths traversing from each NBD Walker B region to each pore loop using the *cnapath* function and the Girvan-Newman algorithm (14) implemented in the Bio3D package (15,16). In the allosteric network, each Cα atom represents a node, and network edges are weighted by $w_{ij} = -\log(|C_{ij}|)$. To remove weakly correlated and physically distant residue pairs, we set $|Cij| \geqslant 0.3$ and the Cα -Cα distance $d_{ij} \leqslant 10$ Å (16). The optimal path corresponds to the shortest distance between the "source" (Walker B) and "sink" (pore loop) residue pairs, whereas suboptimal paths are slightly longer paths, excluding the optimal one.

Similarity between probability density distributions is evaluated using the overlapping coefficient (17,18) is $\mathrm{OC} = \int \min[p_1(\mathrm{x}), p_2(\mathrm{x})]\,\mathrm{dx}$, where $p_i(\mathrm{x}), i = 1, 2,$ are the probability densities to compare.

**Supplemental Figures:**

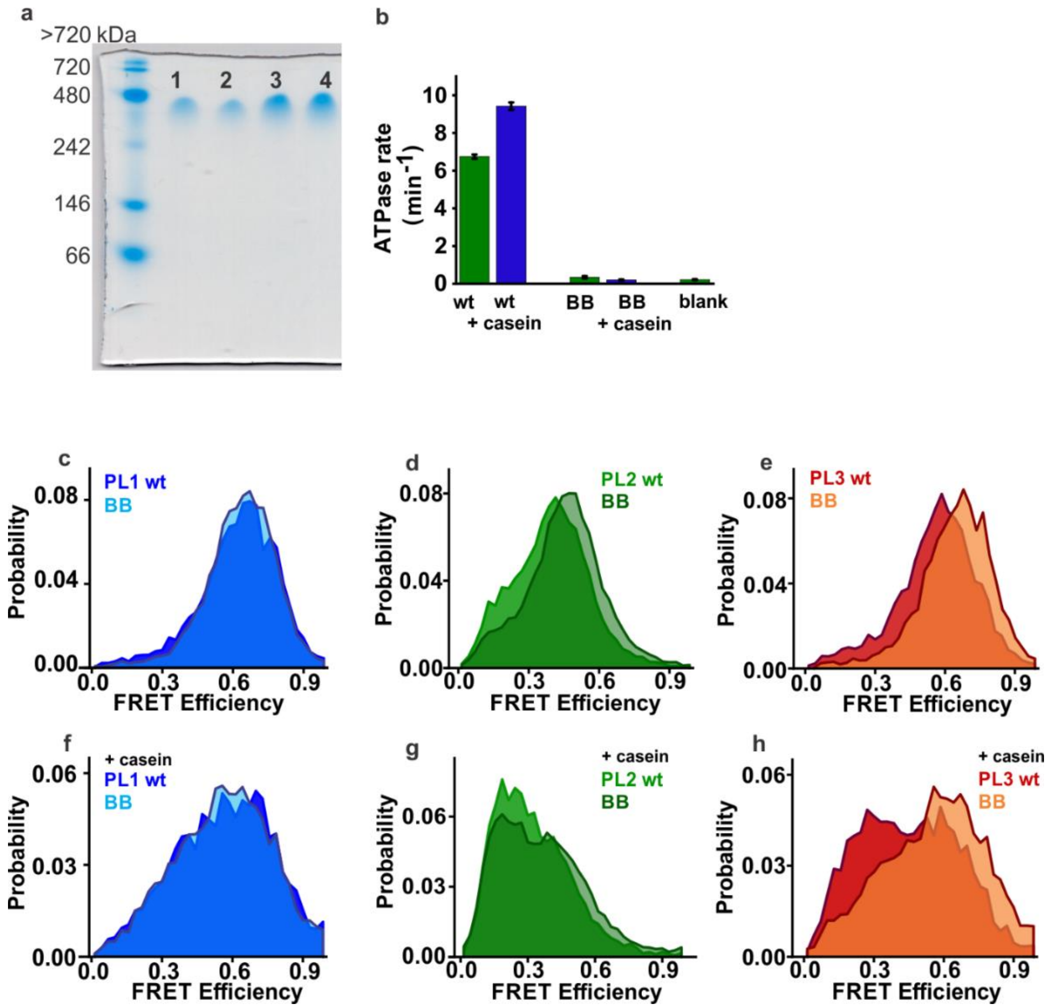

**Figure S1. Double-Walker B (BB) mutants of ClpB show correct assembly but abolished ATP hydrolysis rate and display differential behavior in smFRET experiments. (a)** Native gel (6% acrylamide), stained with Coomassie Blue. Lanes 1, 2 - unmodified dNClpB (referred to as "ClpB" or "wt") and lanes 3, 4 - its double mutant E271A/E668A (numbering as in full-length *TT* ClpB, and denoted as 'BB' here and elsewhere below). Both unmodified wt and BB migrate as a single band in the presence of ATP (run at 4°C, 30 V, 2 mM ATP, 4 mM $Mg^{2+}$) at the molecular weight ~480 kDa (NativeMark$^{TM}$ Unstained Protein Standard, Thermo Fisher Scientific Inc). This is consistent with their homogeneous assembly into dNClpB hexamers (theoretical MW=484.8 kDa). **(b)** Basal and κ-casein (25 μM) stimulated ATPase activities at 25°C (in green and blue, respectively) of wt dNClpB and BB mutant (standard error, n=5). Blank is recorded without added protein. As expected, the BB mutations completely abolish ATP hydrolysis and casein-induced stimulation. **(c)** FRET efficiency histograms of PL1 and double-Walker B construct. Unmodified PL1 ("wt") is in solid blue and PL1 BB ("BB") in semi-transparent cyan. **(d)** PL2 wt is in solid green and PL2 BB in semi-transparent dark-green. **(e)** PL3 wt is in solid red and PL3 BB in semi-transparent orange. **(f)** PL1 wt and PL1 BB with 25 μM κ-casein. Colors as in (a). **(g)** PL2 wt and PL2 BB with 25 μM κ-casein. Colors as in (b). **(h)** PL3 wt and PL3 BB with 25 μM κ-casein. Colors as in (c). Here and elsewhere below, single-molecule measurements were conducted using 1:100 labeled:unlabeled ClpB with 2 mM ATP.

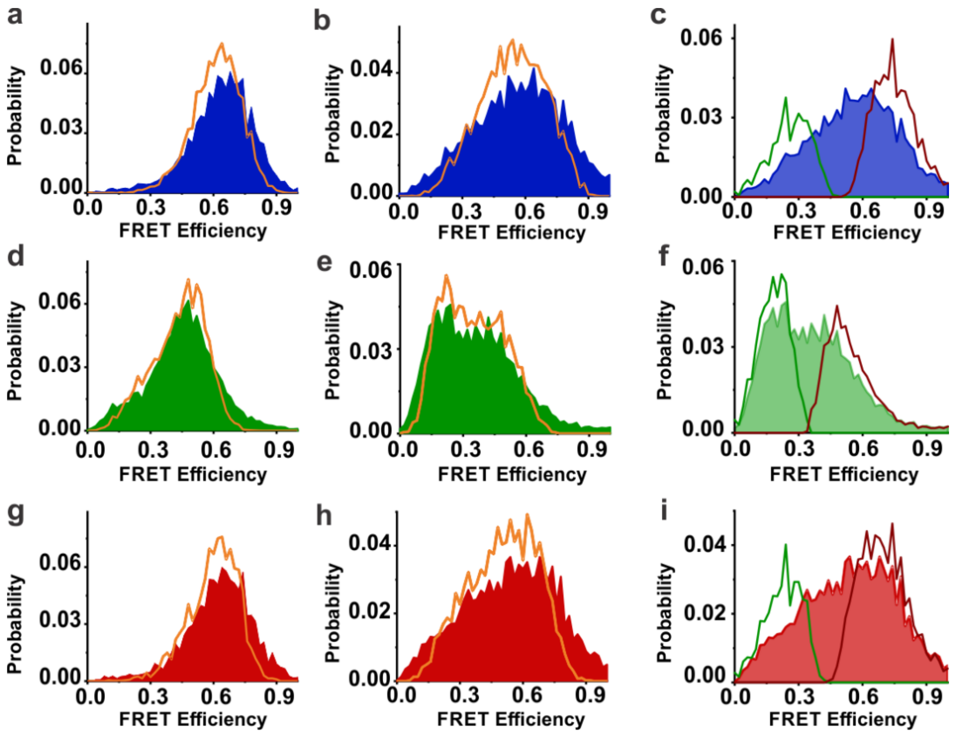

**Figure S2. H²MM analysis of Walker BB mutants.** Recoloring and segmentation analyses of single-molecule FRET data were carried as previously described (1,3,19). See more details under "Supplemental smFRET Data Analysis Details". **(a,b,d,e,g,h)** Recoloring of FRET-efficiency histograms of PL BB data. Recolored histograms are shown as orange curves. **(a)** PL1 BB. **(b)** PL1 BB + κ-casein (25 μM). **(d)** PL2 BB. **(e)** PL2 BB + κ-casein (25 μM). **(g)** PL3 BB. **(h)** PL3 BB + κ-casein (25 μM). In all cases, the quality of recoloring is comparable to that for PLs wt datasets (see our preceding study (1)), even though a fixed-state model is used instead of free model, with the same FRET efficiency values used for different mutants of the same pore-loop type (as detailed in the section "Supplemental smFRET Data Analysis Details" and in Table S1). Histograms are normalized to the sum of events (>7,000 per measurement) and presented with 50 bins. **(c)** Segmentation analysis of FRET efficiency histogram of PL1 BB + κ-casein (25 μM), **(f)** of PL2 with κ-casein (25 μM), **(i)** of PL3 with κ-casein (25 μM). Segmentation analyses clearly show two states, at low FRET efficiency (green curve) and at high FRET efficiency (dark-red curve).

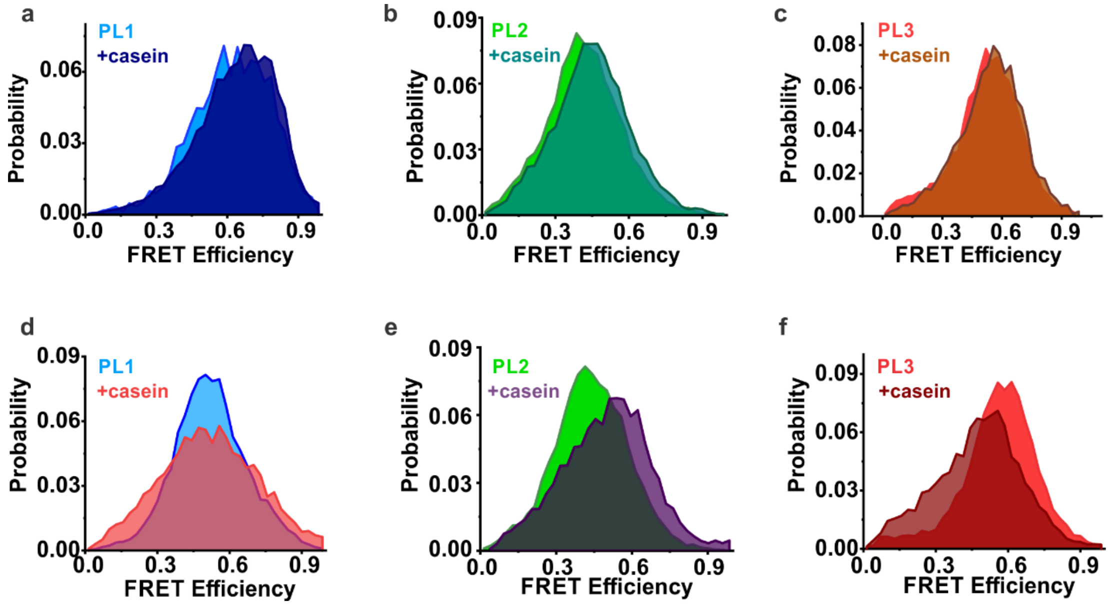

**Figure S3. FRET efficiency histograms from experiments under assembly-disfavoring conditions. (a-c)** Measurements in the presence of 300 mM KCl, without ATP (apo), (25 mM HEPES, 300 mM KCl, 10 mM $MgCl_2$, 0 mM ATP), where disassembly is expected. Addition of κ-casein (25 μM) causes no shift to low FRET efficiency under these conditions. **(d-f)** Measurements in the absence of added ATP and $Mg^{2+}$ (25 mM HEPES, 25 mM KCl, 0 mM $MgCl_2$, 0 mM ATP), where disassembly is also expected. Addition of κ-casein (25 μM) causes a smaller shift to low FRET efficiency in comparison to the measurements in the presence of 2 mM ATP and 10 mM $Mg^{2+}$. Casein-induced shift in FRET efficiency histograms is decreased in all three types of the pore loops, especially under high-salt conditions where hexamers of ClpB are expected to be disassembled. These results suggest that casein-induced shift to low FRET efficiency values occurs only in ATP-bound and assembled ClpB.

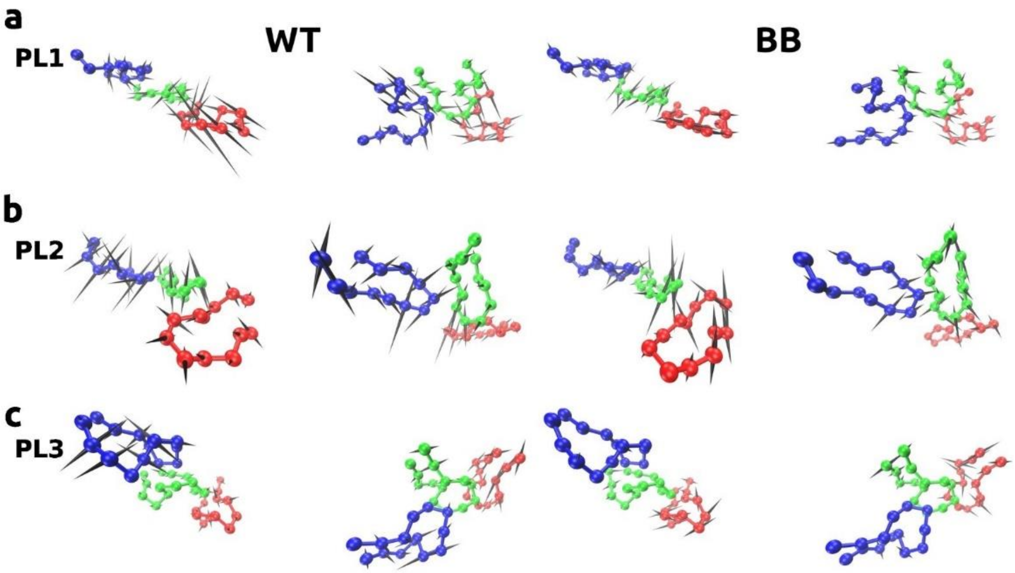

**Figure S4. Pore loop motions associated with PC2 eigenvectors.** Directions of motion of amino acids of (a) PL1 (b) PL2 (c) PL3 in protomers 2-4 (blue, green and red, respectively) are indicated using spikes for the wild-type (left panels) and BB variants (right panels). Top and side views are shown for both the wild-type and BB variants. The amplitudes of motions, indicated by spike lengths, have been scaled by a factor of 2 for clarity.

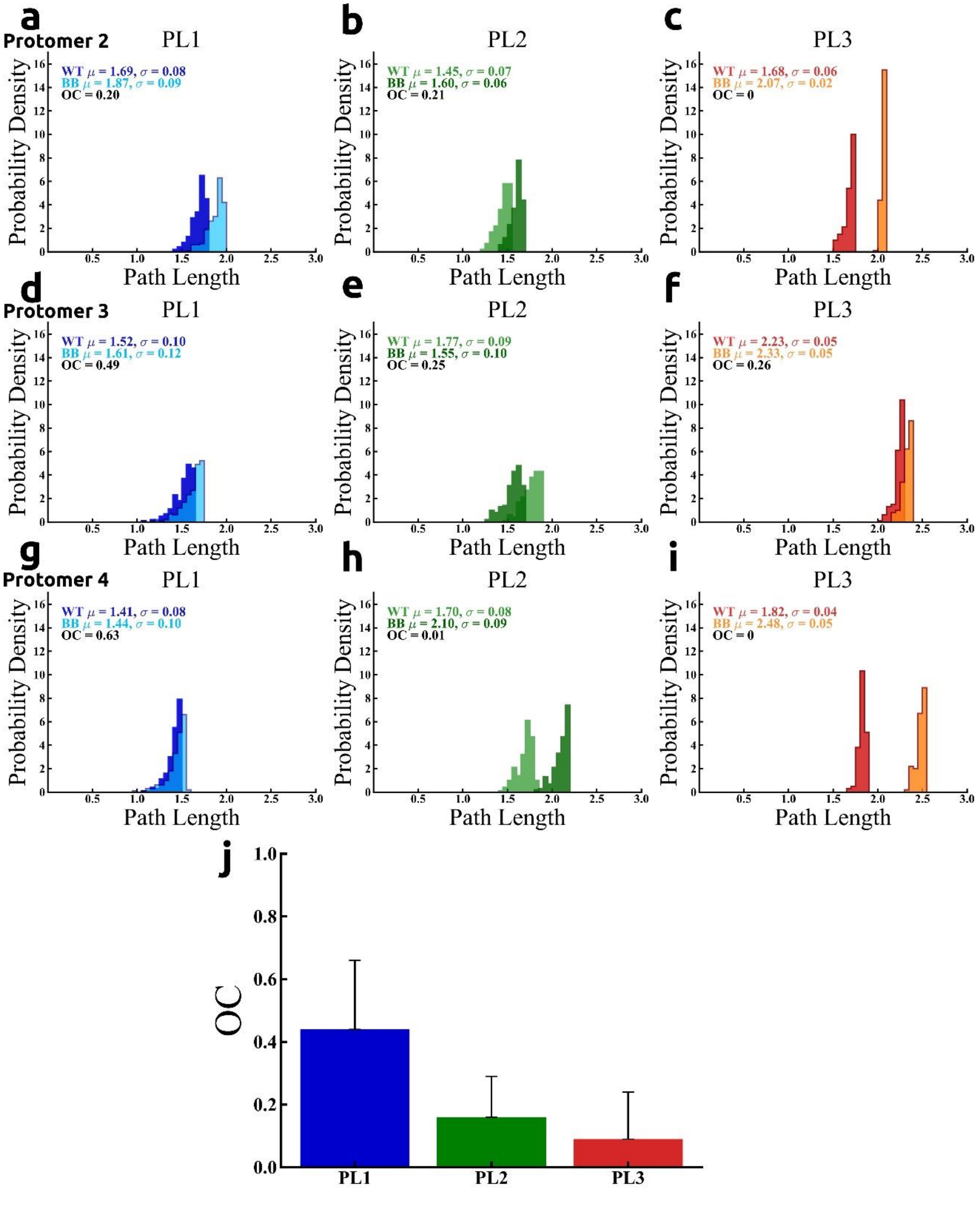

**Figure S5. Path length distributions in allosteric signaling in ClpB configurations.** Probability density distributions of the 200 shortest path lengths are shown for PL1, PL2, and PL3 wt and BB in protomers **(a-c)** 2, **(d-f)** 3, and **(g-i)** 4. **(j)-(k)** Similarity of path length distributions of PL1-3 are quantified using the average and standard deviation (shown as half error bar), over protomers 2-4, of **(j)** the overlap coefficient (OC). The effect of perturbation on allosteric communication is weak in PL1, but strong in PL2 and PL3.

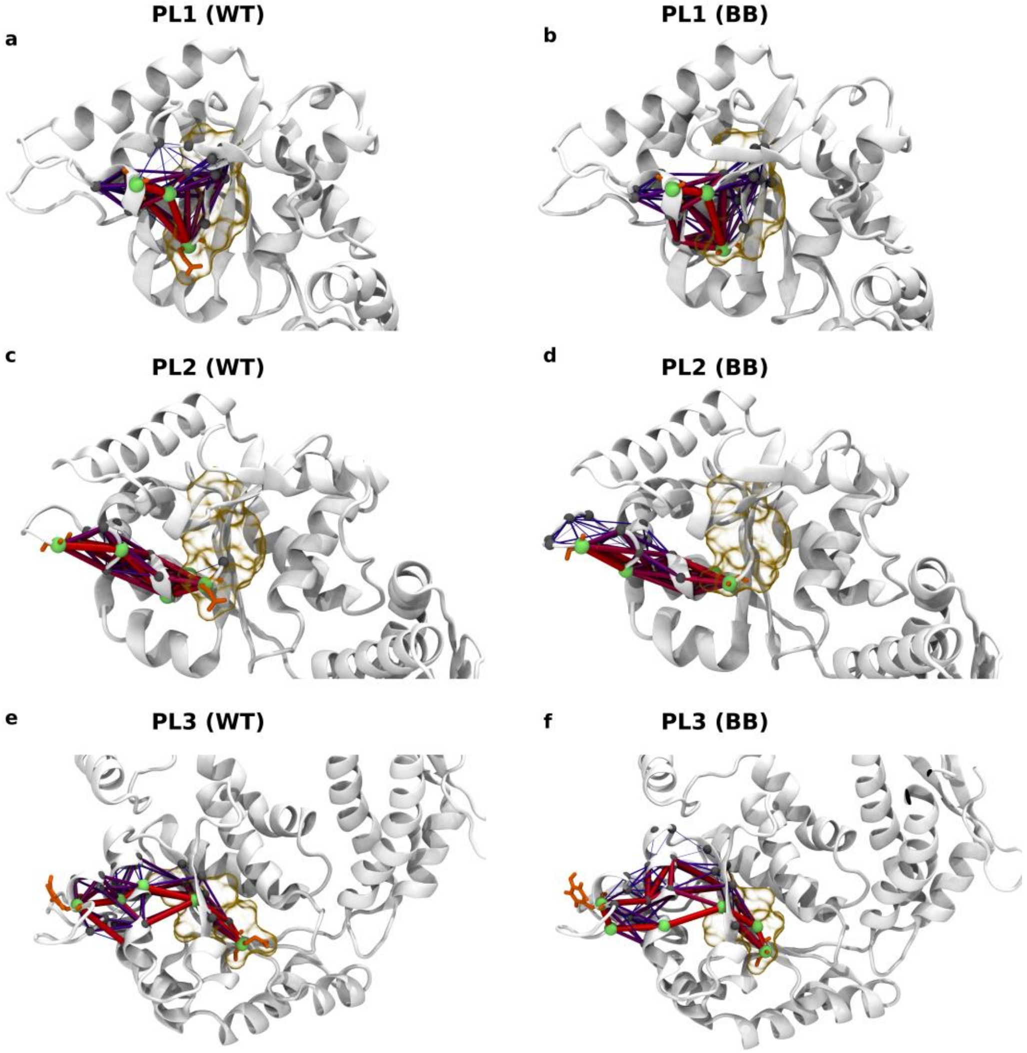

**Figure S6. Optimal and suboptimal paths connecting the Walker B regions and pore loops within a ClpB protomer.** Paths are shown for **(a-b)** PL1 wt and BB; **(c-d)** PL2 wt and BB; **(e-f)** PL3 wt and BB. The pore loop residues labeled in smFRET experiments, A244 in PL1, A289 in PL2, and Y656 in PL3 (numbering for *E. coli* ClpB) are shown in orange using licorice representation. Optimal paths are slightly perturbed by BB mutations. The ensemble of suboptimal paths (purple), which have longer path lengths, is weakly perturbed by BB mutations for PL1 and strongly perturbed for PL2 and PL3. Line thickness is proportional to the strength of the coupling.

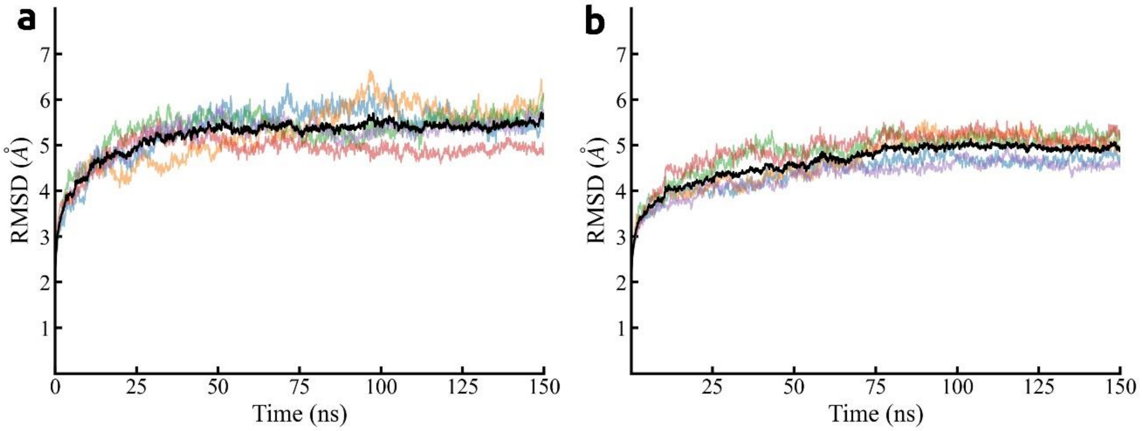

**Figure S7. Equilibration of MD trajectories.** Root-mean-square deviation calculated as the average over five trajectories (black) and for individual trajectories (color-coded) of **(a)** wild-type ClpB and **(b)** BB variant.

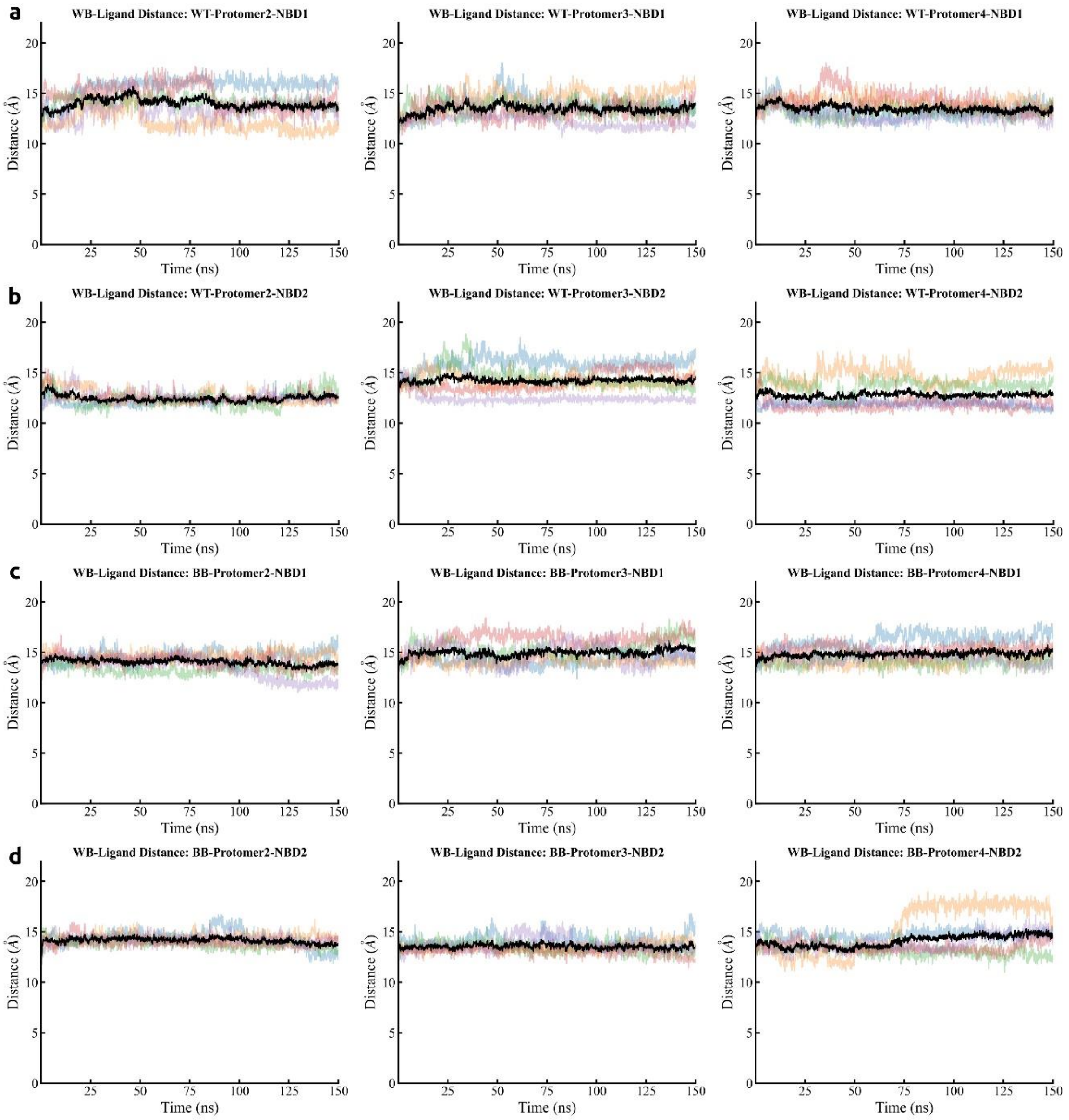

**Figure S8. Nucleotide stability within the binding site.** The distance between the center of mass of nucleotides and the $C_\alpha$ atom of the Walker B mutation location within the same nucleotide binding site in (a) NBD1 and (b) NBD2 of wild-type ClpB and (c) NBD1 and (d) NBD2 of the BB variant. Time series in individual trajectories are indicated using color-coding and averages are indicated in black.

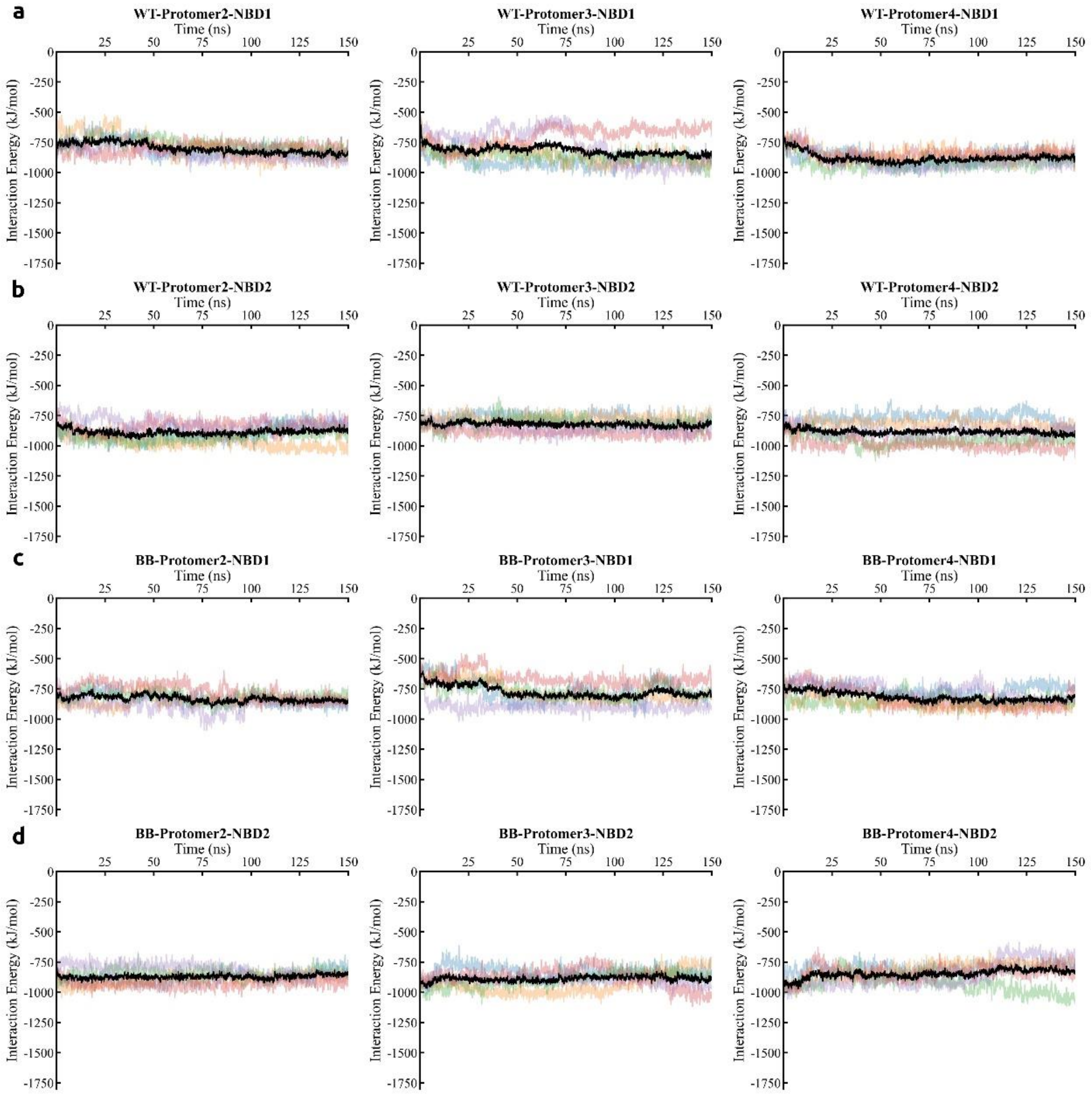

**Figure S9. Interaction between nucleotide and ClpB.** Interaction energy between (a)-(b) wild-type ClpB and nucleotides in (a) NBD1 and (b) NBD2. (c)-(d) BB variant and nucleotides in (c) NBD1 and (d) NBD2. Time series in individual trajectories are indicated using color-coding and averages are indicated in black.

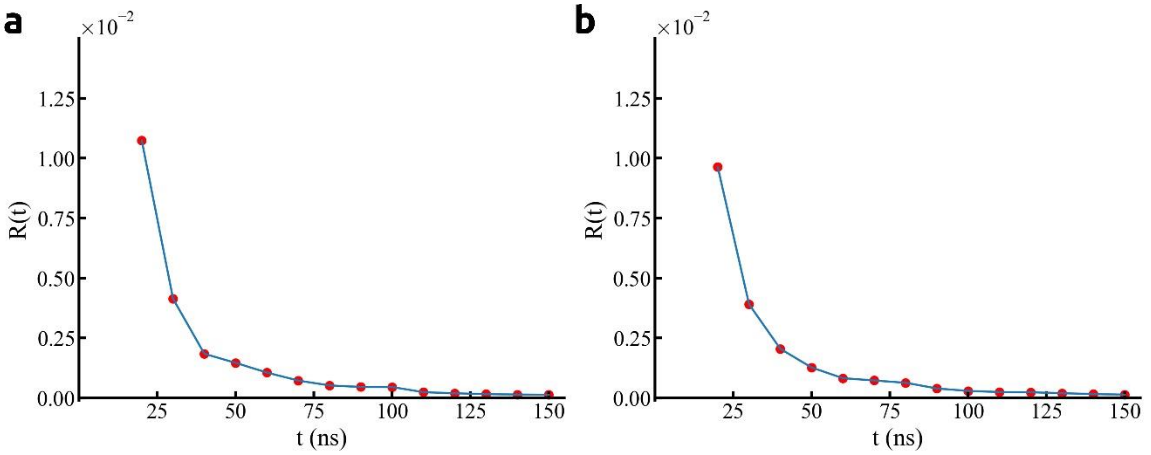

**Figure S10. DCCM convergence.** Mean-square distance computed over multiple trajectories of **(a)** wild-type **(b)** BB variants.

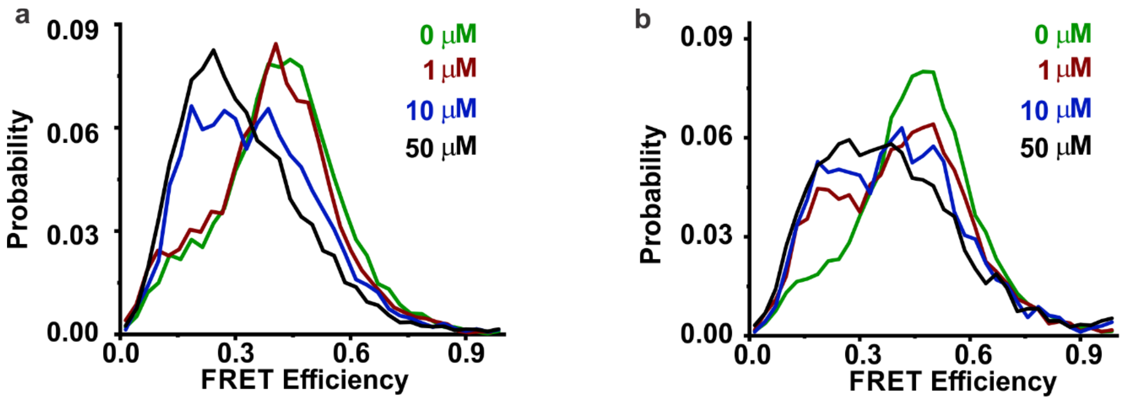

**Figure S11. Representative FRET efficiency histograms from single-molecule κ-casein titration assay. (a)** PL2 wt. **(b)** PL2 BB. Low-FRET state becomes more populated with increasing concentration of κ-casein. This effect is more prominent for the BB mutant, where a strong change is seen already with 1 μM κ-casein.

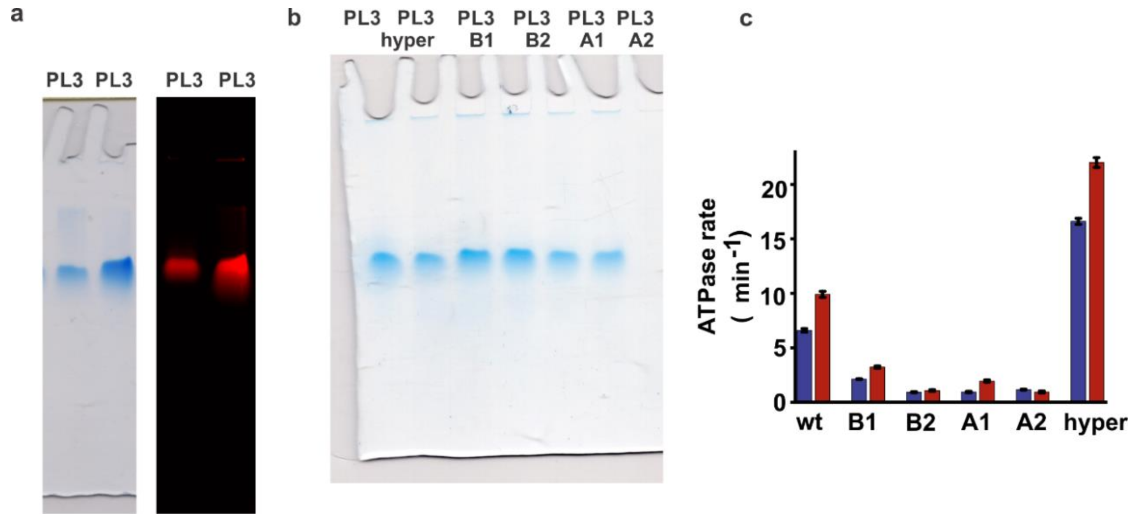

**Figure S12. Single mutants of ClpB display correct assembly but modified ATPase activity.** **(a)** and **(b)** Native PAGE analysis of 1:100 labeled:unlabeled ClpB samples studied in this work. These typically display single bands, similarly to the results for wt dNClpB (see Fig. S1), indicating homogeneously assembled complexes. **(a)** Representative native gel (6% acrylamide) of 1:100 mixed samples of S359C-Y646C (PL3) with wt. Left: stained with Coomassie Blue. Right: same gel as on the left, imaged with a Typhoon scanner at 532 nm confirms incorporation of fluorescently-labelled molecules into the sample following the mixing procedure. Run in the presence of ATP (at 4°C, 30 V, 2 mM ATP, 4 mM $Mg^{2+}$). **(b)** Representative native gel (6% acrylamide, run and stained as in **(a)**) of 1:100 mixed S359C-Y646C (PL3) sample and its single mutants. The single mutations here and elsewhere below are as follows: K347A (‘hyper’), E271A (‘B1’), E668A (‘B2’), K204T (‘A1’) and K601A (‘A2’), with numbering as in the full-length *TT* ClpB. Note that here and elsewhere, the A/B mutations are numbered according to the NBD that bears the mutation (1 or 2). **(c)** ATPase activity and casein-induced enhancement of unlabeled cysteine-less single mutants of ClpB. Background-corrected basal (blue bars) and κ-casein (25 µM) stimulated (red bars) ATPase activities at 25°C of wt (dNClpB) and mutants (standard error, n=5). Hyperactive mutation shows 2.5x higher basal rate than wt, and single Walker A and B are characterized by decreased basal ATPase activity. Most of the mutants show κ-casein-induced activity enhancement.

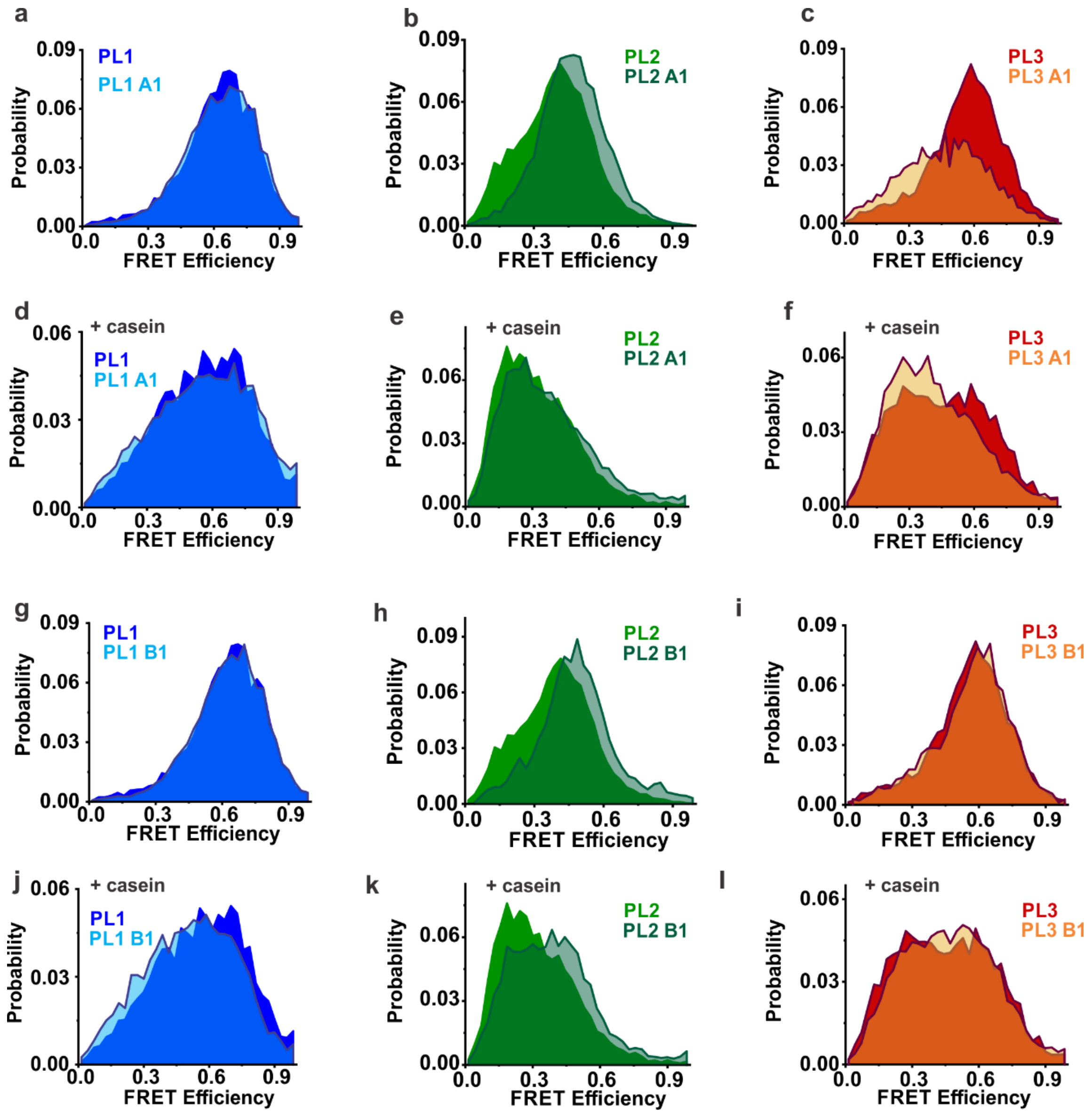

**Figure S13. FRET efficiency histograms of NBD1 mutants.** PL1, PL2 and PL3 without mutations, and Walker A1 (K204T) and Walker B1 (E271A) results. "+ casein" denotes measurements with κ-casein (25 μM). Color scheme as in Fig. S1.

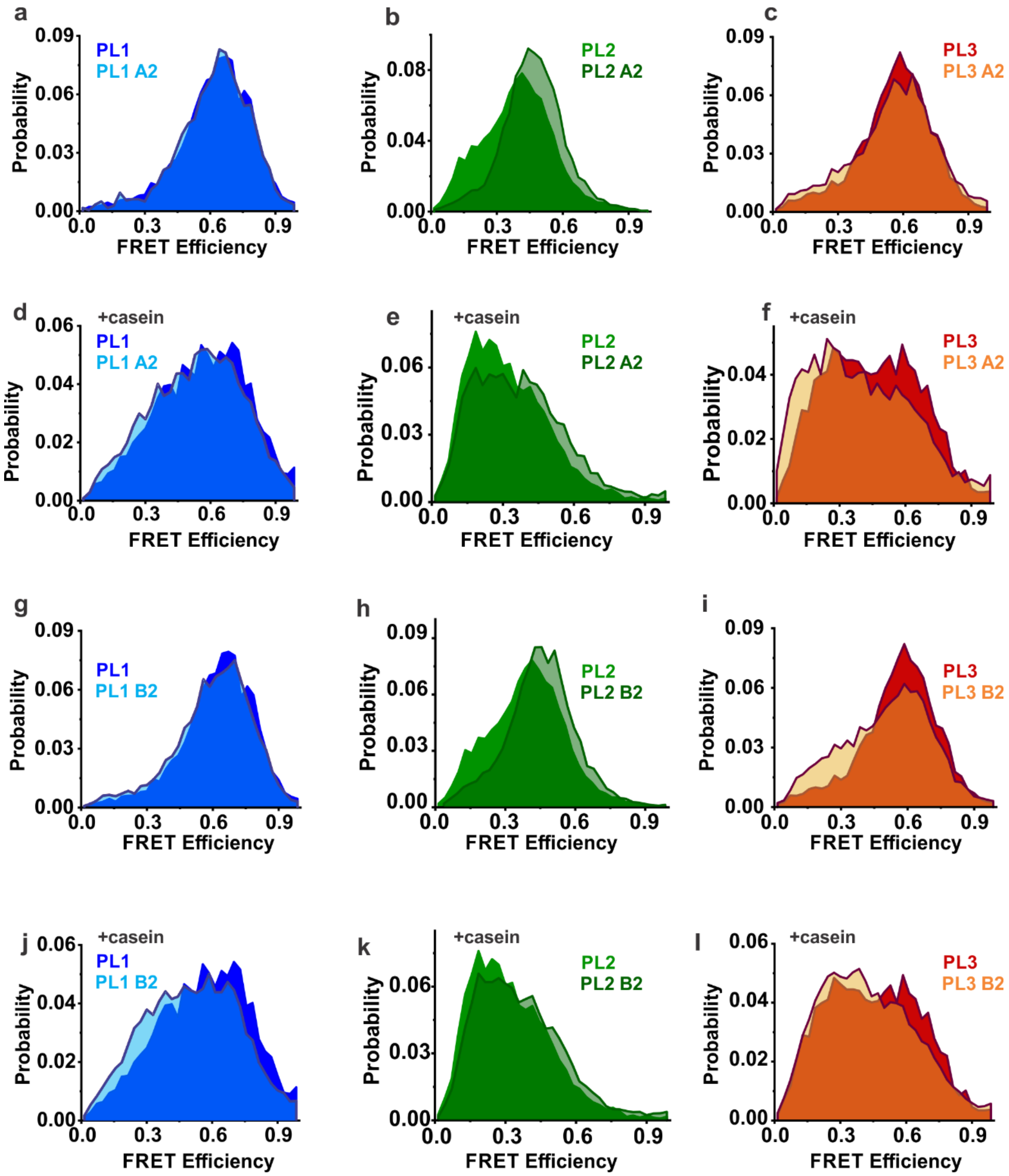

**Figure S14. FRET efficiency histograms of NBD2 mutants.** PL1, PL2 and PL3 without mutations, and Walker A2 (K601T) and Walker B2 (E668A) results. "+ casein" denotes measurements with κ-casein (25 μM). Color scheme as in Fig. S1.

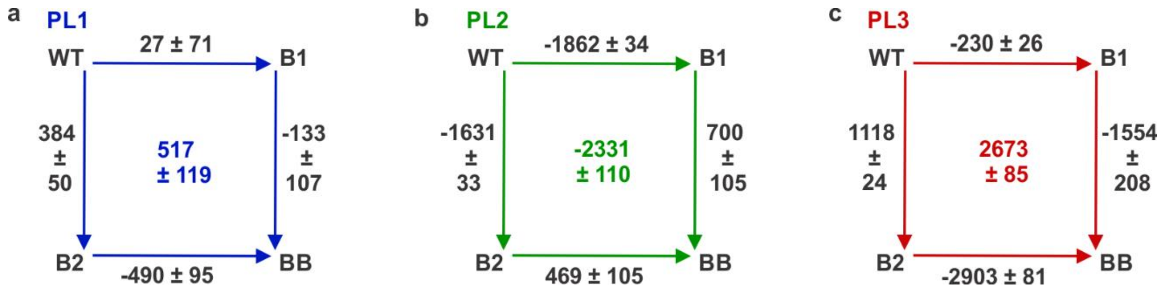

**Figure S15. Double-mutant cycles** (20) for PL1, PL2 and PL3 without κ-casein. The values along the sides are the free energy changes associated with the indicated mutations, $\Delta\Delta G_i$ (in J.mol$^{-1}$), calculated from the H$^2$MM-derived $K_i$s as detailed in Materials and Methods, main text. In these cycles, the $\Delta\Delta G_i$ values along the opposite edges are unequal: $\Delta\Delta G_{i_{WT\rightarrow B1}} \neq \Delta\Delta G_{i_{B2\rightarrow BB}}$ and $\Delta\Delta G_{i_{WT\rightarrow B2}} \neq \Delta\Delta G_{i_{B1\rightarrow BB}}$. This indicates thermodynamic coupling of the effect of the two mutations on pore-loop conformations. The coupling energies for each pore-loop type (Materials and Methods, main text) are shown at the centres of the cycles. Error is from the propagation of the standard error in $K_i$.

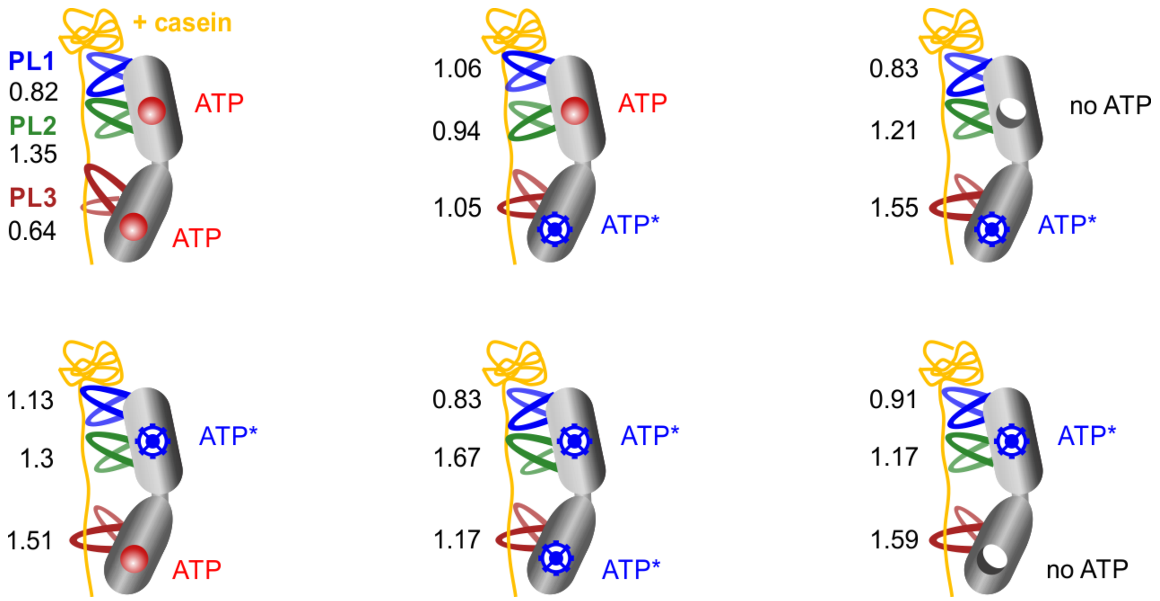

**Figure S16. Summary of all smFRET results suggests an ATP-dependent modulation of pore-loop dynamics.** As in Figure 7 (main text), states of ClpB pore loops are schematically shown, with bound κ-casein in yellow, and PL1, PL2 and PL3 in blue, green and dark-red, respectively. The ATPase states of the NBDs are depicted as follows: red circles are bound ATP molecules not undergoing hydrolysis (ATP arrested state), blue wheels – bound ATP undergoing hydrolysis, which corresponds to a mixture of ATP/ADP, and empty circles – unbound or apo state. To note, these apo states are likely to be very transient under our experimental conditions (excess ATP at 2 mM). Nevertheless, they represent important states within the ATPase cycle (essential precursors for the ATP re-binding to occur), and their presence in ClpB was previously experimentally captured by cryo-EM (21). As in Figure 7 (main text), PLs can favor either an up or a down conformation, or visit both states with almost equal probability, depending on the ATPase state of the NBDs. The size of the PLs in the scheme reflects their state occupancy based on the $H^2MM$ analysis, and the numbers are $H^2MM$-derived equilibrium coefficients, $K_i$s.

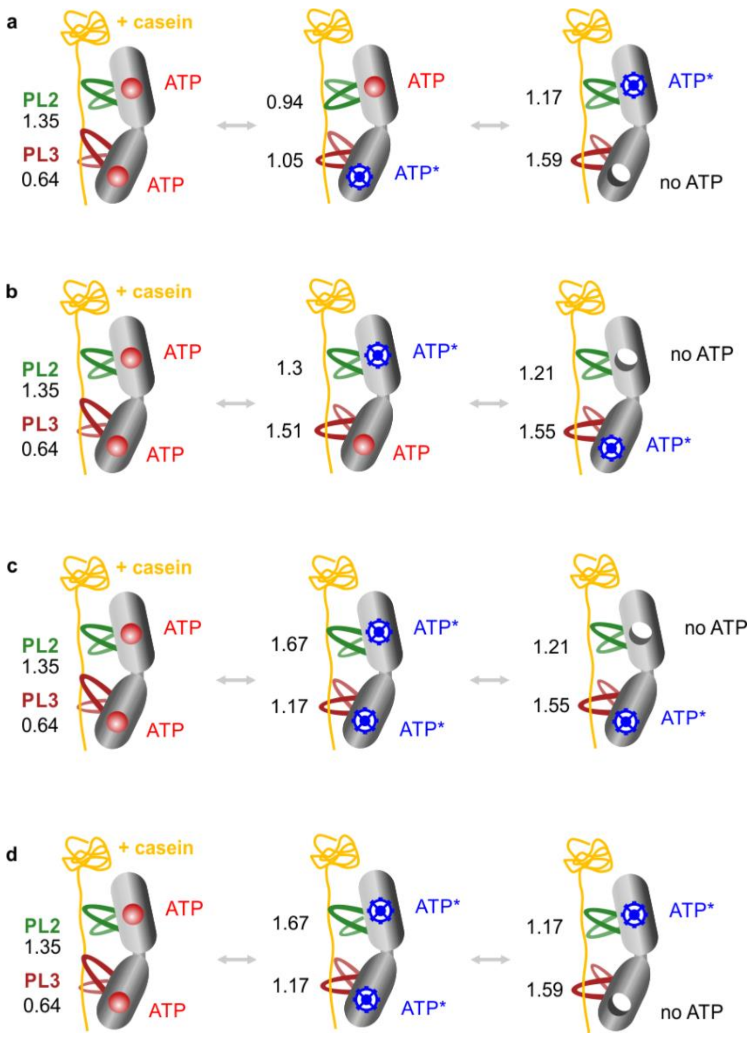

**Figure S17. Examples of potential routes leading to substrate-protein translocation by ClpB. (a-d)** These routes can be considered under a few assumptions: 1) The order of events is based on the ATP hydrolysis reaction (ATP binds → ATP hydrolysis event → nucleotide release results in apo state → ATP re-binds). 2) There is no preference for the order of ATP binding/hydrolysis happening in NBD1 or in NBD2. 3) A translocation event occurs upon a change in the equilibrium position of PL3 from up to down state, happening within a single protomer of ClpB. During this transition, PL3 is either pulling on the substrate-protein, or is acting as a pawl to guide its movement, as discussed in the main text. Here, there is no requirement for any specific conformations of PL2 relative to PL3. To point out, the functioning protomer still needs to be embedded into a hexamer to ensure correct ATP binding and hydrolysis. There is no information on the states of neighboring protomers because only intra-protomer dynamics are being measured in our assays, and therefore substrate-protein translocation driven by the movements of two or more protomers within a hexamer cannot be inferred. For simplicity, PL1 is omitted since this pore-loop type did not show strong ATP-dependent modulations. The size of the PLs on the scheme reflects their state occupancy from the $H^2MM$ analysis, and the numbers are $H^2MM$-derived equilibrium coefficients, $K_i$s.

**Supplemental Movies:**

**Movie SM1.** Motions associated with the principal component PC1 of PL1 loops in wild-type ClpB (left) and in the BB variant (right).

**Movie SM2.** Motions associated with the principal component PC1 of PL2 loops in wild-type ClpB (left) and in the BB variant (right).

**Movie SM3.** Motions associated with the principal component PC1 of PL3 loops in wild-type ClpB (left) and in the BB variant (right).

**Supplemental Data Summary Tables:**

**Table S1. FRET efficiency values of the two pore loop states as obtained from the global $H^2$MM analysis of wild-type pore-loop constructs, measured with or without κ-casein** (1)**.**

| **Pore loop** | **FRET value** State 1 | **FRET value** State 2 |
|---|---|---|
| **PL1** | 0.240 | 0.767 |
| **PL2** | 0.182 | 0.536 |
| **PL3** | 0.212 | 0.699 |

**Table S2. Cumulative overlap of the top 2 principal components of pore loops in the BB variant over the top 10 principal components of the corresponding pore loops in wild-type ClpB.**

| Cumulative Overlap | BB-PL1-PC1 | BB-PL1-PC2 | BB-PL2-PC1 | BB-PL2-PC2 | BB-PL3-PC1 | BB-PL3-PC2 |
|---|---|---|---|---|---|---|
| wt-PC1 | 0.8451 | 0.0782 | 0.1004 | 0.0926 | 0.1158 | 0.0790 |
| wt-PC2 | 0.8452 | 0.5416 | 0.2260 | 0.5088 | 0.2556 | 0.0865 |
| wt-PC3 | 0.8777 | 0.6099 | 0.2281 | 0.5589 | 0.2677 | 0.1110 |
| wt-PC4 | 0.8778 | 0.6113 | 0.3076 | 0.5792 | 0.5208 | 0.1522 |
| wt-PC5 | 0.8913 | 0.6275 | 0.3653 | 0.5817 | 0.5208 | 0.3335 |
| wt-PC6 | 0.8915 | 0.7394 | 0.4602 | 0.7048 | 0.5225 | 0.3859 |
| wt-PC7 | 0.9641 | 0.7397 | 0.6203 | 0.7070 | 0.6305 | 0.4611 |
| wt-PC8 | 0.9694 | 0.7560 | 0.6642 | 0.7070 | 0.6766 | 0.5630 |
| wt-PC9 | 0.9700 | 0.7801 | 0.8026 | 0.7421 | 0.6922 | 0.6519 |
| wt-PC10 | 0.9734 | 0.7824 | 0.8172 | 0.7678 | 0.7410 | 0.6881 |

**Table S3: Optimal paths between Walker B sites and pore loops in NBD1, PL1 and PL2, in wild type ClpB and BB mutants, derived from the MD simulations of *E. coli* ClpB.**

| Protomer | PL1 | |
|---|---|---|
| | Wild-type | BB |
| 2 | A279 → M283 → M242 → G243 → A244 | A279 → M242 → G243 → A244 |
| 3 | A279 → A241 → A244 | A279 → M242 → G243 → A244 |
| 4 | A279 → A241 → A244 | A279 → A241 → A244 |

| Protomer | PL2 | |
|---|---|---|
| | Wild-type | BB |
| 2 | A279 → M283 → A286 → K288 → A289 | A279 → L280 → V284 → G285 → K288 → A289 |
| 3 | A279 → M283 → A287 → K288 → A289 | A279 → L280 → V284 → K288 → A289 |
| 4 | A279 → L280 → H281 → A286 → A289 | A279 → L280 → V284 → G285 → A289 |

**Table S4: Optimal paths between Walker B sites and pore loops in NBD2, PL3, in wild type ClpB and BB mutants, derived from the MD simulations of *E. coli* ClpB.**

| Protomer | PL3 | |
|---|---|---|
| | Wild-type | BB |
| 2 | A678 → D677 → I632 → L646 → V647 → G648 → Y656 | A678 → M716 → I715 → V714 → V713 → T712 → F709 → V707 → D701 → Y656 |
| 3 | A678 → D677 → I632 → L646 → A649 → Y656 | A678 → D677 → I632 → L646 → A647 → Y656 |
| 4 | A678 → I632 → Y661 → G660 → Y656 | A678 → D677 → I632 → R645 → A649 → Y656 |

**Table S5. Substrate κ-casein binding constants to individual pore-loop mutants, derived from $H^2$MM analysis of single-molecule κ-casein titration measurements.**

| Sample | $K_d$ (μM) | Error (from fit)* | $F_{max}$ | Error (from fit) |
|---|---|---|---|---|
| **PL1 wt** | 2.9 | 0.5 | 0.56 | 0.02 |
| **PL1 BB** | 2.0 | 0.4 | 0.57 | 0.04 |
| **PL2 wt** | 11.3 | 3.0 | 0.83 | 0.10 |
| **PL2 BB** | 0.3 | 0.1 | 1.03 | 0.03 |
| **PL3 wt** | 0.9 | 0.4 | 0.74 | 0.06 |
| **PL3 BB** | 1.6 | 0.7 | 0.97 | 0.09 |

*Data, presented in Fig. 5 (main text) were fitted to a simple binding isotherm $y = 1 + \frac{\text{Fmax} \times [\text{casein}]}{(K_d + [casein])}$

where [casein] is κ-casein concentration (μM) and $F_{max}$ is the maximum P1(+ casein)/P1(- casein).

**Tables S6: H$^2$MM Analysis Outputs.** Average state populations, equilibrium constants, $K_i$ and state-to-state transition rates, $k^i$, (in $s^{-1}$) for each pore loop type $(i)$, compared to dwell time analysis.

**PL1: WT and double mutants.**

| Pore loop sampe+2 mM ATP | State Populations | | $K_i$ | $k^i_{12}$ | | $k^i_{21}$ | |
|---|---|---|---|---|---|---|---|
| | State 1 | State 2 | | H$^2$MM | Dwell time | H$^2$MM | Dwell time |
| **PL1** | 0.32 ± 0.01 | 0.68 ± 0.01 | 0.46 ± 0.02 | 44370 ± 1750 | 42360 ± 1700 | 20640 ± 1780 | 23630 ± 470 |
| **PL1 + κ-casein** | 0.45 ± 0.01 | 0.55 ± 0.01 | 0.83 ± 0.02 | 17670 ± 2230 | 21490 ± 1190 | 14730 ± 1720 | 19160 ± 1070 |
| **PL1 BB** | 0.31 ± 0.02 | 0.69 ± 0.02 | 0.45 ± 0.05 | 47580 ± 5990 | 42700 ± 4130 | 22460 ± 1900 | 23160 ± 770 |
| **PL1 BB + κ-casein** | 0.45 ± 0.01 | 0.55 ± 0.01 | 0.82 ± 0.02 | 21350 ± 810 | 22900 ± 470 | 17470 ± 560 | 20460 ± 250 |

The errors (standard deviation) were calculated from at least three measurements. Here and elsewhere below, "+ κ-casein" is with 25 µM κ-casein.

**PL2: WT and double mutants.**

| Pore loop sampe+2 mM ATP | State Populations | | $K_i$ | $k^i_{12}$ | | $k^i_{21}$ | |
|---|---|---|---|---|---|---|---|
| | State 1 | State 2 | | H$^2$MM | Dwell time | H$^2$MM | Dwell time |
| **PL2** | 0.44 ± 0.01 | 0.56 ± 0.01 | 0.79 ± 0.02 | 11230 ± 880 | 13090 ± 90 | 8890 ± 510 | 11100 ± 290 |
| **PL2 + κ-casein** | 0.62 ± 0.01 | 0.38 ± 0.01 | 1.67 ± 0.04 | 5950 ± 280 | 9500 ± 310 | 9910 ± 370 | 12180 ± 200 |
| **PL2 BB** | 0.33 ± 0.02 | 0.67 ± 0.02 | 0.49 ± 0.05 | 10700 ± 1380 | 13540 ± 180 | 5260 ± 150 | 9000 ± 80 |
| **PL2 BB + κ-casein** | 0.57 ± 0.02 | 0.43 ± 0.02 | 1.35 ± 0.1 | 4680 ± 190 | 6210 ± 520 | 6230 ± 200 | 7440 ± 40 |

The errors (standard deviation) were calculated from at least two measurements.

**PL3: WT and double mutants.**

| Pore loop sampe+2 mM ATP | State Populations | | $K_i$ | $k^i_{12}$ | | $k^i_{21}$ | |
|---|---|---|---|---|---|---|---|
| | State 1 | State 2 | | H$^2$MM | Dwell time | H$^2$MM | Dwell time |
| **PL3** | 0.34 ± 0.01 | 0.66 ± 0.01 | 0.50 ± 0.02 | 28300 ± 4100 | 29730 ± 3780 | 14250 ± 2020 | 18430 ± 1760 |
| **PL3 + κ-casein** | 0.54 ± 0.01 | 0.46 ± 0.01 | 1.17 ± 0.06 | 9650 ± 240 | 13470 ± 560 | 11330 ± 250 | 13680 ± 80 |
| **PL3 BB** | 0.20 ± 0.01 | 0.80 ± 0.01 | 0.24 ± 0.02 | 33890 ± 3700 | 26540 ± 1170 | 8280 ± 1290 | 11330 ± 980 |
| **PL3 BB + κ-casein** | 0.39 ± 0.01 | 0.61 ± 0.01 | 0.64 ± 0.04 | 13440 ± 580 | 15090 ± 240 | 8530 ± 110 | 12080 ± 710 |

The errors (standard deviation) were calculated from at least two measurements.

**<u>PL1:</u> Single-NBD mutants.**

| Pore loop sampe+2 mM ATP | State Populations | | $K_i$ | $k^i_{12}$ | | $k^i_{21}$ | |
|---|---|---|---|---|---|---|---|
| | State 1 | State 2 | | H$^2$MM | Dwell time | H$^2$MM | Dwell time |
| **PL1 A1** | 0.31 ± 0.01 | 0.69 ± 0.01 | 0.45 ± 0.03 | 46490 ± 1330 | 41280 ± 340 | 20760 ± 1960 | 23480 ± 1000 |
| **PL1 A1 + κ-casein** | 0.45 ± 0.01 | 0.55 ± 0.01 | 0.83 ± 0.05 | 15140 ± 920 | 20560 ± 770 | 12540 ± 70 | 18530 ± 80 |
| **PL1 A2** | 0.32 ± 0.01 | 0.68 ± 0.01 | 0.47 ± 0.02 | 41750 ± 290 | 42090 ± 400 | 19550 ± 600 | 24020 ± 420 |
| **PL1 A2 + κ-casein** | 0.48 ± 0.01 | 0.52 ± 0.01 | 0.91 ± 0.05 | 19300 ± 1980 | 26040 ± 1550 | 17570 ± 850 | 24100 ± 230 |
| **PL1 B1** | 0.32 ± 0.01 | 0.68 ± 0.01 | 0.47 ± 0.03 | 48460 ± 830 | 42090 ± 750 | 22770 ± 1930 | 24240 ± 1430 |
| **PL1 B1 + κ-casein** | 0.51 ± 0.01 | 0.49 ± 0.01 | 1.06 ± 0.02 | 14640 ± 4640 | 22530 ± 3130 | 15450 ± 4580 | 23220 ± 2680 |
| **PL1 B2** | 0.35 ± 0.01 | 0.65 ± 0.01 | 0.54 ± 0.01 | 41090 ± 7610 | 41100 ± 4230 | 22260 ± 3630 | 25700 ± 1950 |
| **PL1 B2 + κ-casein** | 0.53 ± 0.01 | 0.47 ± 0.01 | 1.13 ± 0.07 | 15010 ± 550 | 22860 ± 800 | 16900 ± 380 | 23820 ± 540 |

The errors (standard deviation) were calculated from at least two measurements.

**<u>PL1:</u> Other conditions**

| Pore loop sampe+2 mM ATP | State Populations | | $K_i$ | $k^i_{12}$ | | $k^i_{21}$ | |
|---|---|---|---|---|---|---|---|
| | State 1 | State 2 | | H$^2$MM | Dwell time | H$^2$MM | Dwell time |
| **PL1 hyper (K347A)** | 0.28 ± 0.01 | 0.72 ± 0.01 | 0.39 ± 0.01 | 46920 ± 90 | 43540 ± 1930 | 18320 ± 730 | 22060 ± 1170 |
| **PL1 hyper (K347A) + κ-casein** | 0.41 ± 0.01 | 0.59 ± 0.01 | 0.69 ± 0.03 | 19050 ± 3330 | 24120 ± 4780 | 13050 ± 1650 | 19330 ± 2760 |
| **PL1 apo 300 mM KCl** | 0.35 ± 0.03 | 0.65 ± 0.03 | 0.53 ± 0.08 | 45330 ± 7670 | 43630 ± 4050 | 23780 ± 410 | 27140 ± 330 |
| **PL1 apo 300 mM KCl + κ-casein** | 0.32 ± 0.03 | 0.68 ± 0.03 | 0.47 ± 0.06 | 40140 ± 4320 | 39120 ± 3990 | 18710 ± 220 | 23170 ± 360 |
| **PL1 apo no $Mg^{2+}$** | 0.49 ± 0.02 | 0.51 ± 0.02 | 0.98 ± 0.09 | 49570 ± 110 | 39400 ± 430 | 48590 ± 4530 | 38700 ± 2290 |
| **PL1 apo no $Mg^{2+}$ + κ-casein** | 0.52 ± 0.01 | 0.48 ± 0.01 | 1.10 ± 0.02 | 21480 ± 1300 | 22480 ± 1130 | 23600 ± 1930 | 23490 ± 1074 |

The errors (standard deviation) were calculated from at least two measurements.

**<u>PL2:</u> Single-NBD mutants.**

| Pore loop sampe+2 mM ATP | State Populations | | $K_i$ | $k^i_{12}$ | | $k^i_{21}$ | |
|---|---|---|---|---|---|---|---|
| | State 1 | State 2 | | H²MM | Dwell time | H²MM | Dwell time |
| **PL2 A1** | 0.26 ± 0.003 | 0.74 ± 0.003 | 0.35 ± 0.01 | 23920 ± 1730 | 19450 ± 860 | 8390 ± 480 | 10620 ± 310 |
| **PL2 A1 + κ-casein** | 0.55 ± 0.02 | 0.45 ± 0.02 | 1.21 ± 0.11 | 6410 ± 580 | 9390 ± 500 | 7730 ± 30 | 11080 ± 50 |
| **PL2 A2** | 0.25 ± 0.005 | 0.75 ± 0.005 | 0.33 ± 0.01 | 25150 ± 790 | 18110 ± 370 | 8360 ± 60 | 10570 ± 120 |
| **PL2 A2 + κ-casein** | 0.54 ± 0.02 | 0.46 ± 0.02 | 1.17 ± 0.09 | 5870 ± 50 | 9720 ± 30 | 6850 ± 440 | 11070 ± 70 |
| **PL2 B1** | 0.27 ± 0.01 | 0.73 ± 0.01 | 0.37 ± 0.01 | 16210 ± 2490 | 16430 ± 310 | 6000 ± 770 | 9990 ± 450 |
| **PL2 B1 + κ-casein** | 0.48 ± 0.03 | 0.52 ± 0.03 | 0.94 ± 0.1 | 8970 ± 780 | 11570 ± 560 | 8340 ± 270 | 11410 ± 130 |
| **PL2 B2** | 0.29 ± 0.01 | 0.71 ± 0.01 | 0.41 ± 0.01 | 18980 ± 800 | 15130 ± 180 | 7740 ± 140 | 10110 ± 10 |
| **PL2 B2 + κ-casein** | 0.56 ± 0.02 | 0.44 ± 0.02 | 1.3 ± 0.1 | 6580 ± 790 | 10710 ± 860 | 8340 ± 390 | 13610 ± 2330 |

The errors (standard deviation) were calculated from at least two measurements.

**<u>PL2:</u> Other conditions**

| Pore loop sampe+2 mM ATP | State Populations | | $K_i$ | $k^i_{12}$ | | $k^i_{21}$ | |
|---|---|---|---|---|---|---|---|
| | State 1 | State 2 | | H²MM | Dwell time | H²MM | Dwell time |
| **PL2 hyper (K347A)** | 0.22 ± 0.01 | 0.78 ± 0.01 | 0.28 ± 0.01 | 28200 ± 1270 | 22660 ± 400 | 8030 ± 80 | 11050 ± 670 |
| **PL2 hyper (K347A) + κ-casein** | 0.46 ± 0.005 | 0.54 ± 0.005 | 0.87 ± 0.02 | 8460 ± 170 | 10970 ± 150 | 7350 ± 290 | 11160 ± 400 |
| **PL2 apo 300 mM KCl** | 0.37 ± 0.01 | 0.63 ± 0.01 | 0.59 ± 0.02 | 16480 ± 730 | 9740 ± 70 | 18240 ± 800 | 12210 ± 5 |
| **PL2 apo 300 mM KCl + κ-casein** | 0.33 ± 0.01 | 0.67 ± 0.01 | 0.50 ± 0.02 | 17500 ± 620 | 8720 ± 50 | 16590 ± 240 | 10800 ± 140 |
| **PL2 apo no $Mg^{2+}$** | 0.33 ± 0.005 | 0.67 ± 0.005 | 0.49 ± 0.01 | 18300 ± 1200 | 16900 ± 630 | 8950 ± 390 | 10640 ± 290 |
| **PL2 apo no $Mg^{2+}$ + κ-casein** | 0.26 ± 0.01 | 0.74 ± 0.01 | 0.36 ± 0.01 | 9280 ± 460 | 10900 ± 1100 | 3300 ± 280 | 7400 ± 440 |

The errors (standard deviation) were calculated from at least two measurements.

**<u>PL3:</u> Single-NBD mutants.**

| Pore loop sampe+2 mM ATP | State Populations | | $K_i$ | $k^i_{12}$ | | $k^i_{21}$ | |
|---|---|---|---|---|---|---|---|
| | State 1 | State 2 | | H²MM | Dwell time | H²MM | Dwell time |
| **PL3 A1** | 0.48 ± 0.01 | 0.52 ± 0.01 | 0.91 ± 0.04 | 19780 ± 1940 | 16840 ± 3 | 17950 ± 1020 | 16060 ± 350 |
| **PL3 A1 + κ-casein** | 0.61 ± 0.01 | 0.39 ± 0.01 | 1.55 ± 0.07 | 11860 ± 180 | 11890 ± 210 | 18370 ± 550 | 15620 ± 230 |
| **PL3 A2** | 0.36 ± 0.01 | 0.64 ± 0.01 | 0.57 ± 0.03 | 19770 ± 420 | 17400 ± 690 | 11200 ± 430 | 12210 ± 500 |
| **PL3 A2 + κ-casein** | 0.61 ± 0.02 | 0.39 ± 0.02 | 1.59 ± 0.15 | 5500 ± 180 | 9200 ± 90 | 8750 ± 540 | 11080 ± 140 |
| **PL3 B1** | 0.31 ± 0.01 | 0.69 ± 0.01 | 0.46 ± 0.01 | 26780 ± 3700 | 21850 ± 1310 | 12240 ± 1310 | 13030 ± 170 |
| **PL3 B1 + κ-casein** | 0.51 ± 0.01 | 0.49 ± 0.01 | 1.05 ± 0.05 | 15030 ± 400 | 13170 ± 630 | 15820 ± 370 | 13730 ± 400 |
| **PL3 B2** | 0.44 ± 0.01 | 0.56 ± 0.01 | 0.79 ± 0.03 | 13570 ± 790 | 14320 ± 370 | 10780 ± 280 | 12060 ± 50 |
| **PL3 B2 + κ-casein** | 0.60 ± 0.01 | 0.40 ± 0.01 | 1.51 ± 0.04 | 9840 ± 50 | 11250 ± 60 | 14860 ± 310 | 14000 ± 210 |

The errors (standard deviation) were calculated from at least two measurements.

**<u>PL3:</u> Other conditions**

| Pore loop sampe+2 mM ATP | State Populations | | $K_i$ | $k^i_{12}$ | | $k^i_{21}$ | |
|---|---|---|---|---|---|---|---|
| | State 1 | State 2 | | H²MM | Dwell time | H²MM | Dwell time |
| **PL3 hyper (K347A)** | 0.25 ± 0.02 | 0.75 ± 0.02 | 0.33 ± 0.03 | 45400 ± 1670 | 28420 ± 380 | 14730 ± 900 | 13590 ± 40 |
| **PL3 hyper (K347A) + κ-casein** | 0.41 ± 0.004 | 0.59 ± 0.004 | 0.69 ± 0.01 | 17830 ± 1430 | 14280 ± 390 | 12290 ± 780 | 13350 ± 440 |
| **PL3 apo 300 mM KCl** | 0.38 ± 0.03 | 0.62 ± 0.03 | 0.60 ± 0.07 | 35970 ± 5420 | 35850 ± 2760 | 21500 ± 660 | 23800 ± 10 |
| **PL3 apo 300 mM KCl + κ-casein** | 0.36 ± 0.01 | 0.64 ± 0.01 | 0.56 ± 0.02 | 37070 ± 840 | 35770 ± 190 | 20770 ± 1030 | 23510 ± 490 |
| **PL3 apo no $Mg^{2+}$** | 0.32 ± 0.02 | 0.68 ± 0.02 | 0.48 ± 0.05 | 41520 ± 1710 | 27070 ± 380 | 19910 ± 2830 | 14910 ± 90 |
| **PL3 apo no $Mg^{2+}$ + κ-casein** | 0.49 ± 0.01 | 0.51 ± 0.01 | 0.96 ± 0.02 | 18670 ± 380 | 16260 ± 2840 | 17880 ± 770 | 15440 ± 2330 |

The errors (standard deviation) were calculated from at least two measurements.
